# Triploid Pacific oysters exhibit stress response dysregulation and elevated mortality following marine heatwaves

**DOI:** 10.1101/2023.03.02.530828

**Authors:** Matthew N. George, Olivia Cattau, Mollie Middleton, Delaney Lawson, Brent Vadopalas, Mackenzie Gavery, Steven Roberts

## Abstract

Polyploidy has been shown to negatively impact environmental stress tolerance, resulting in increased susceptibility to extreme climate events such as marine heatwaves (MHWs). In this study, we used the response of the Pacific oyster *Crassostrea gigas* to MHWs as a model system to identify key ploidy-specific differences in the physiological and transcriptomic response of oysters to environmental stress. In this study, adult diploid (2n) and triploid (3n) oysters were exposed to elevated seawater temperature (single stressor; 30°C), elevated temperature followed by acute desiccation stress (multiple stressor; 30°C + emersion at an aerial temperature of 44°C for 4h), or a control (17°C) within a hatchery setting. Oyster mortality rate was elevated within stress treatments with respect to the control and was significantly higher in triploids than diploids following multiple stress exposure (36.4% vs. 14.8%). Triploids within the multiple stressor treatment exhibited signs of energetic limitation, including metabolic depression, a significant reduction in ctenidium Na^+^/K^+^ ATPase activity, and the dysregulated expression of key genes associated with heat tolerance, the inhibition of apoptosis, and mitochondrial function. Functional enrichment analysis of ploidy-specific gene sets identified that biological processes associated with metabolism, stress tolerance, and immune function were overrepresented within triploids across stress treatments. Our results demonstrate that triploidy impacts the transcriptional regulation of key metabolic processes that underly the environmental stress response of Pacific oysters, resulting in downstream shifts in physiological tolerance limits that may be detrimental to survival. The impact of chromosome set manipulation on the climate resilience of marine populations has important implications for the adaptability of marine populations and domestic food security within future climate scenarios, especially as triploidy induction becomes an increasingly popular tool to elicit reproductive control across a wide range of marine organisms used within marine aquaculture.

## 1. Introduction

Chromosome set duplication (polyploidy) occurs when there is total nondisjunction of chromosomes during mitosis or meiosis, resulting in cells or organisms with one or more additional homologous chromosome sets. Polyploid animals generally have larger cells, lower cell membrane surface area to volume ratios, slower cell division rates, and achieve larger sizes as adults (‘polyploid gigantism’) than diploid (2n) conspecifics (Gregory & Mable, 2005). Cell size differences have meaningful consequences for metabolism; smaller cells require more energy to maintain ionic gradients between the cytoplasm and their surroundings (Rolfe & Brown, 1997; Szarski, 1983) and have faster oxygen and nutrient diffusion rates than larger cells (Subczynski et al., 1989). As a result, polyploids have been reported to have lower mass-specific metabolic rates than diploids (Kozłowski et al., 2003; Starostová et al., 2009), are more susceptible to oxygen limitation under elevated temperature (Hermaniuk et al., 2021), and perform better within cold environments (Atkins & Benfey, 2008; Sambraus et al., 2017).

While phenotypic differences between polyploids and their diploid progenitors have been described within a variety of taxa, a mechanistic understanding of the relationship between polyploidy and stress tolerance remains elusive (Soltis et al., 2016). Polyploidy has been associated with performance benefits (i.e., heterosis) and consequences within stressful environments (Van de Peer et al., 2021). Potential benefits of polyploidy include enhanced genetic and phenotypic diversity (Fox et al., 2020), resilience to DNA damage (Schoenfelder & Fox, 2015), and the suppression of tumors (Bielski et al., 2018); potential consequences include epigenome and genome instability (Storchova, 2014), gene loss (Moghe & Shiu, 2014), energetic costs associated with enhanced gene silencing (Doyle & Coate, 2019; Mittelsten Scheid et al., 1996), and transcriptome changes (related to changes in gene dosage) that alter the function of regulatory networks (Coate et al., 2016; Comai, 2005).

A useful model system with which to investigate the impact of polyploidy on stress tolerance is oyster aquaculture. Techniques to induce triploidy (3n) in oysters were first developed in the 1980s and are now widely adopted by industry (Downing & Allen, 1987; Guo et al., 2009; Nell, 2002; Yamamoto et al., 1988). Oyster aquaculture leverages triploidy to induce reproductive impairment in an attempt to avoid the seasonal, energy-intensive cycle of gametogenesis that can result in enhanced mortality within the summer months (a phenomenon known as ‘summer mortality’; Perdue et al., 1981; Samain & McCombie, 2008). By partitioning energy towards somatic growth, triploid oysters display enhanced growth and meat weight, decreased time to market, and greater marketability year round (Allen & Downing, 1986, 1991; Dégremont et al., 2012; Normand et al., 2008; Shpigel et al., 1992). Early investigations of the ability of triploidy to compensate for summer mortality in France were promising, with triploid oysters exhibiting lower mortality rates (Gagnaire et al., 2006; Samain, 2011), presumably due to enhanced energetic investment in growth and maintenance processes (Jouaux et al., 2013). However, in practice, triploids can experience enhanced mortality when compared with diploids grown under the same field conditions (known as ‘triploid mortality’). Triploid mortalities have been observed in France (Houssin et al., 2019), as well as the Eastern (Guévélou et al., 2019; Matt, 2018; Matt et al., 2020), Southern (Wadsworth, 2018; Wadsworth et al., 2019) and Western (Tim Morris, Pacific Seafood; Paul Taylor, Taylor Shellfish Co.; Kurt Grinnell, Jamestown S’Klallam Tribe, personal communications) coasts of the United States, resulting in substantial economic losses. While the selective mortality of triploid oysters has been linked to environmental variability at specific sites (*C. virginica:* Bodenstein et al., 2021; Guévélou et al., 2019; *C. gigas:* B. Eudeline, Taylor Shellfish, personal communication), it remains unclear whether ploidy manipulation, and the physiological and transcriptomic perturbations that result, enhance or decrease the risk of summer mortality.

The summer mortality rates of marine organisms are expected to increase along with the likelihood and intensity of marine heatwaves (MHWs). MHWs are prolonged extreme oceanic high-temperature events that have far reaching ecological and economic consequences, with individual events resulting in direct economic losses of $800 million and more than $3.1 billion in indirect losses for multiple consecutive years (Smith et al., 2021). For intertidal organisms, MHWs can present multiple abiotic stressors in rapid succession that can work either synergistically or antagonistically to impact physiology (Breitburg & Riedel, 2005). A poignant demonstration of this phenomenon was observed when the Pacific Northwestern region of the United States experienced a MHW that coincided with some of the lowest daytime low tides of the year during the summer of 2021 (Philip et al., 2021; White et al., 2022). Exposure to elevated seawater temperatures followed by desiccation stress through emersion over three tidal cycles resulted in the death of an estimated one billion shellfish within the coastal and inland waters of Washington State and British Columbia and substantial losses for commercial and tribal growers (Cecco, 2021; Raymond et al., 2022).

The impact of polyploidy on the susceptibility to summer mortality following MHWs remains unclear. Differences in energy storage could benefit polyploids within multiple stressor scenarios, while changes in the genetic architecture that underlies important stress response pathways could alter physiological tolerance limits (Holbrook et al., 2020). Using MHW and oyster aquaculture as a model, we conducted a hatchery experiment to identify key ploidy-specific differences in the stress response of diploid and triploid *C. gigas* using an energy-limited tolerance to stress framework (Sokolova, 2013; Sokolova et al., 2012). Two-year-old commercially produced oysters were exposed to either a single stressor (elevated seawater temperature, 30C), multiple stressors in succession (elevated seawater temperature, 30°C + emersion for 4h @ 44°C), or a control and observed for mortality for up to 30 days. Differences in the physiological and transcriptomic response of oysters to single and multiple environmental stressors across ploidy were identified across multiple levels of biological organization by monitoring changes in standard metabolic rate, metabolic enzyme activity rates, and gene expression to identify key biological processes associated with stress tolerance impacted by polyploidy.

## 2. Methods

Five hundred diploid (2n) and triploid (3n) Pacific oysters (*Crassostrea gigas*) with varied parentage were obtained from commercial partners. Triploidy was induced through the exposure of larvae to a thermal shock during meiosis I (Yamamoto et al., 1988). Adult oysters were received in March, 2021 and transferred to the Jamestown S’Klallam Point Whitney Shellfish Hatchery located on Dabob Bay, Washington (47°45”N 122°50’W). Upon receipt, the ploidy of 15 individuals from each group was determined using a CyFlow™ Ploidy Analyzer (Sysmex America Inc., Lincolnshire, IL, USA); remaining oysters were individually labeled by affixing numbered wire tags to the right shell valve using cyanoacrylate. The maximum shell length, width, and height of each oyster was recorded using vernier calipers to the nearest mm; measurements were taken at the beginning and end of experiments. Shell volume was estimated from measurements using the equation of a generalized ellipsoid. To determine the baseline condition of a subset of oysters upon arrival at the hatchery, gonadal sections were sampled for histological examination and dry tissue weight was measured by drying excised whole-body tissue at 60°C for 72 h. Dry tissue weight and reproductive condition (see section 2.2.) was determined for remaining oysters following tissue sampling or after the conclusion of experiments. In the event that dry weight could not be determined for an individual due to mortality or other constraints, dry weight was estimated from shell volume (see Figure S1).

### 2.1. Experimental Design

All oysters were allowed to acclimate to hatchery conditions (T ~ 17°C, salinity ~ 27 ppt, pH ~ 8; see table 1) for twenty days (day −30 to −10) within 100 L tanks with flowing seawater from Dabob Bay. Throughout the experiment, a peristaltic pump was used to mix live algae into seawater lines to achieve a constant cell density of 3 × 10^5^ cells ml^−1^ within experimental tanks. Oysters were fed a mixed algal diet consisting of an equal proportion of *Tetraselmis* spp., *Pavlova* spp., and *Chaetoceros* spp. Following acclimation, diploid and triploid oysters were haphazardly split into three treatment groups: control, single stressor, or multiple stressor (see Figure 1A for timeline); treatment groups were as follows: diploid control (2n-C), diploid single stressor (2n-SS), diploid multiple stressor (2n-MS), triploid control (3n-C), triploid single stressor (3n-SS), and triploid multiple stressor (3n-MS). The starting sample size within each treatment was 112 oysters. Oysters within each treatment were placed in triplicate within PVC silos (76 mm diameter) with netting (3.2 mm mesh) affixed to the bottom, which were themselves placed within individually capped PVC sleeves (101 mm diameter) to separate the effluent of each cluster. A pseudo-replicated design was used to prevent the mortality of an oyster within a given treatment from impacting others outside of its respective cohort. Each silo was supplied seawater at a constant rate of 100 ml min^−1^ and had a turnover rate of 78 min.

**Table 1.** Seawater temperature (°C), salinity (ppt), and pH (NBS scale) within each treatment (± SD).

| Treatment | Temperature | Salinity | pH |
| --- | --- | --- | --- |
| Acclimation | 17.3 ± 0.8<br>(15.6 – 18.9) | 27.5 ± 0.9<br>(26.0 – 29.6) | 8.10 ± 0.08<br>(7.85 – 8.20) |
| Control | 17.1 ± 0.8<br>(14.7 – 18.9) | 28.2 ± 1.7<br>(25.0 – 31.3) | 7.93 ± 0.05<br>(7.81 – 8.01) |
| Single stressor | 30.2 ± 0.5<br>(27.4 – 30.9) | 28.9 ± 1.3<br>(26.2 – 31.6) | 7.97 ± 0.04<br>(7.90 – 8.05) |
| Multi-stressor | 30.4 ± 0.4<br>(28.3 – 30.9) | 28.8 ± 1.0<br>(26.5 – 31.3) | 8.04 ± 0.11<br>(7.90 – 8.23) |

**Figure 1.**
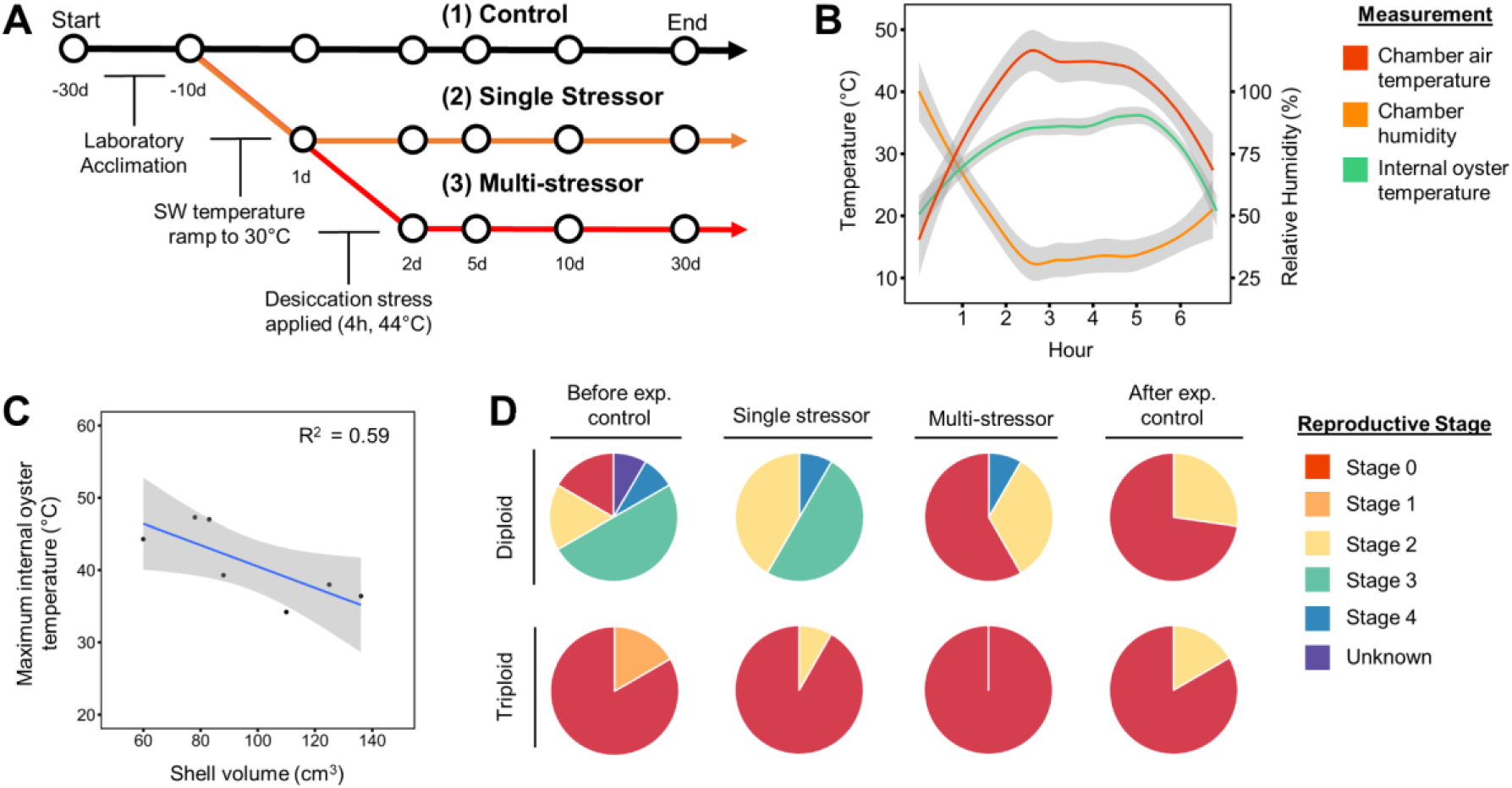
(A) Experimental design including a timeline of when stressors were applied. (B) Desiccation chamber air temperature, humidity, and internal oyster temperature during multiple stressor treatment on day 2. (C) Relationship between shell volume and the maximum internal (meat) temperature experienced by oysters within desiccation chambers. (D) Proportion of oysters within each ploidy at each reproductive stage upon receipt at the hatchery (before experiment control), after single and multiple stress exposure, and within the control treatment after 30 days (after experiment control); twenty-five oysters per ploidy were sampled for reproductive condition per time point.

The temperature (± 0.1°C), salinity (± 0.1), and pH (± 0.01) of each treatment tank was continuously monitored using an A3 Apex Controller System (Neptune Systems, Morgan Hill, CA, USA). The accuracy of each monitored parameter, in addition to the dissolved oxygen concentration (± 0.1 mg L^−1^) within each silo to check for hypoxic conditions, was manually measured every 3-5 days using a HI98199 Multiparameter meter (Hanna Instruments, Woonsocket, RI, USA). Seawater temperature within the control treatment was allowed to match that of the water entering the hatchery, controlled to a maximum setpoint of 17°C within the header tank by an CY-5-CWCT chiller to avoid temperature spikes (Aqualogic, San Diego, CA, USA). Starting at day −10, the temperature within the single and multiple stressor treatments was increased at a rate of 2°C d^−1^ until a setpoint of 30°C was reached (hereafter designated day 0, start of stress treatment); seawater was heated at this rate to approximate the conditions experienced during the lead up to a marine heatwave (Philip et al., 2021). Experimental tanks were heated by passing seawater through two inline Aqualogic Optima Compact Plus Heaters; individual setpoints within each tank were maintained dynamically through the control of three 800-watt titanium rod heaters by a ramp/soak temperature controller utilizing PID control (Auber Instruments, Alpharetta, GA, USA).

On day 2, oysters within the multiple stressor treatment were removed from seawater and placed within 40 L custom-built desiccation chambers for 6 h (see Figure S3). The air temperature within chambers was controlled to a setpoint of 44°C through the dynamic control of three ceramic heat lamps by a temperature controller utilizing PID control. The humidity of each chamber was allowed to fluctuate with temperature. The internal temperature of a subset of oysters was monitored during desiccation trials; oyster tissue temperature was measured in real time by drilling a 1.5 mm hole into the top shell valve of each oyster, inserting a K-type thermocouple into the stomach, and sealing the hole with putty (see Figure S3). While in desiccation chambers, air temperature was held at approximately 44°C for 4 h, with an hour before and after exposure of ramp. Following desiccation, oysters were returned to 30°C seawater. Oysters within the single and multiple stressor treatments were held within 30°C seawater for a total of 30 days. Throughout the experiment, the mortality and metabolic rate of individually labeled oysters within each treatment were repeatedly monitored. Tissue from the ctenidia were also sampled from a subset of oysters preceding and following the application of stress treatments and saved for later enzymatic and transcriptomic analysis. More information regarding the method and sampling regime employed for each metric can be found below.

### 2.2. Reproductive Condition

Oyster reproductive condition (n=15 per ploidy, per treatment, per time point) was determined for a subset of individuals within the control, single stressor, and multiple stressor treatments through sampling and histological analysis of gonad tissue sections. Tissue was sampled from oysters within the control treatment upon receipt at the Point Whitney Hatchery (day −30) and after the conclusion of the experimental trials (day 30). Tissue was sampled from oysters within stressor treatments the day after experimental conditions were applied (single stressor: day 1; multiple stressor: day 2). Sampled gonad tissue was placed within a histology cassette and fixed using the PAXgene tissue fixative and Stabilizer system (Qiagen, Hilden, Germany). Samples were embedded in paraffin, stained with hematoxylin and eosin, sectioned, and imaged at 4x, 10x, and 40x magnification. Images of diploid and triploid gonad sections were assigned to reproductive stages in accordance with ploidy-specific metrics as outlined by (Ezgeta Balic et al., 2020) and (Matt & Allen, 2021), respectively. Gonad was assessed as either resting (stage 0), within an early (stage 1) or late growth stage (stage 2), mature (stage 3), spawning (stage 4), or unknown.

### 2.3. Survival Analysis

Oysters within each treatment group (n=88 per treatment) were checked for mortality every 1-3 days over the course of experiments. The survival R package was used to compute survival curves using the Kaplan-Meier estimator from mortality data (Therneau et al., 2022). The impact of ploidy and stress treatment on survival curves was assessed with the log-rank test using a significance cut off of 0.05.

### 2.4. Standard Metabolic Rate

Whole oyster oxygen consumption rate was measured as a proxy for standard metabolic rate (SMR), at either 17 °C (control) or 30°C (treatment). The SMR of 40 labeled oysters within the control, single stressor, and multiple stressor treatments were taken during the acclimation period (day −15 to −10), followed by repeated measurements of each individual at day 1, 2, 6, and 10 following initial stress exposure. The oxygen uptake rate of individual oysters was determined by closed-system respirometry wherein animals were placed in 500 ml plastic chambers filled with air-saturated, UV-sterilized, and 0.2 um filtered seawater. Chambers were partially submerged within a recirculating water bath controlled to a temperature setpoint. For each measurement, a single oyster was placed within a chamber and allowed to recover from handling stress for at least 15 minutes; chambers were then closed, shielded from light, and the decline in O_2_ concentration (mg O_2_) was recorded for 1 h. Trials in which the oxygen saturation declined below 60% within a chamber were discarded. During each set of trials, the oxygen concentration of a blank chamber with an empty oyster shell was also recorded as an internal control. The mass-specific standard metabolic rate (SMR, mg O2 h^−1^ g^−1^) for each individual at a given time point and temperature was calculated using the following equation:

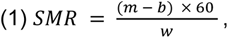

where *m* is the oxygen uptake rate of the linear portion of the oxygen concentration curve (mg min^−1^), *b* is mean oxygen uptake rate of the replicate blanks from that sampling day (mg min^−1^), and *w* is oyster dry tissue weight (g). The oxygen consumption rate of blank chambers as a function of oyster shell size is provided in Figure S2.

### 2.5. Na^+^/K^+^ ATPase

Tissue was collected from diploid and triploid oysters (n=12 per ploidy, per treatment, per time point) within the control, single stressor, and multiple stressor treatments on days −10, 1, 2, 5, 10, and 30. Upon sampling, ctenidia were flash frozen in SEI buffer (250 mM sucrose, 10 mM Na_2_EDTA, 50 mM imidazole; Zaugg, 1982) and stored at −80°C until analysis. The Na^+^/K^+^ ATPase activity of frozen samples was measured within six months according to the method of (McCormick, 1993) with modifications as described by (Bianchini & Wood, 2003) to improve assay performance for *C. gigas*. Briefly, two salt solutions (assay mix A & B) were used during analysis. The ATP concentration within assay mix A and B were decreased to 0.35 mM and the ouabain concentration in assay mix B was increased to 1 mM. Kinetic assays were run at 35°C for ten minutes on a SpectraMax 190 plate reader (Molecular Devices, San Jose, California).

### 2.6. Citrate Synthase

Ctenidium was collected from diploid and triploid oysters (n=12 per ploidy, per treatment, per time point) within the control and single stressor treatments on day 1 and the multiple stressor treatment on day 2, flash frozen, and stored at −80°C for later analysis. Citrate synthase enzyme activity was determined using the Abcam CS Assay Kit (ab239712, Abcam PLC, Cambridge, UK). Briefly, 10-20 mg (wet weight) of ctenidium was homogenized in 350 uL of ice-cold CS Assay Buffer. The optical density of the supernatant in the presence of a developer and substrate (43 uL buffer, 5 uL developer, and 2 uL substrate) was measured in triplicate and averaged at 405 nm for 45-60 minutes at 25°C on a Wallac Victor 2 1420 Multilabel Counter Plate Reader (Perkin Elmer, Waltham, MA, USA). Citrate synthase activity was standardized against the protein concentration of each sample using the Bovine Serum Albumin Assay (500-0201, Bio-Rad Laboratories, Inc., Hercules, CA, USA) and reported as nmol min^−1^ mg^−1^ protein after comparison with a glutathione (GSH) standard.

### 2.7. Differential gene expression (DEG) analysis

Ctenidium was collected from diploid and triploid oysters within the control, single stressor, and multiple stressor treatments on day 1 and 2, flash frozen, and stored at −80°C. Total RNA was extracted from 20 mg tissue samples homogenized in 1 ml of Trizol reagent (RNAzol® RT; Molecular Research Center Inc, Cincinnati, OH). Homogenate was purified using the Direct-zol® RNA Purification Kit (Zymo Research, Irvine, CA) and genomic DNA was removed using the Turbo DNase Kit (Invitrogen, Waltham, MA). RNA was quantified using a Qubit 3.0 Fluorometer (Life Technologies, Carlsbad, CA). RNA integrity was confirmed using the RNA 6000 pico assay on an Agilent 2100 Bioanalyzer (Santa Clara, CA). Twelve biological replicates from each treatment per ploidy (N = 72) were submitted for sequencing at the Genomic Sequencing and Analysis Facility at the University of Texas at Austin. Library preparation was performed using the QuantSeq 3’ mRNA-Seq protocol (v.015UG009V0251, Lexogen, Vienna, Austria), also known as Tag-Seq. Tag-Seq generates cDNA from the 3’ end of mRNA strands with only one fragment per mRNA transcript; this approach avoids many of the computational issues commonly encountered during RNA-seq and the resulting sequence tags are considered direct proxies of gene expression levels. Additionally, tag counts are not influenced by gene length, making it ideal for differential gene expression (DGE) analysis (Asmann et al., 2009; Hong et al., 2011).

Bioinformatic workflows were performed using bash v.4.4.20 (www.gnu.org/software/bash/) and R v4.1.0 (www.r-project.org/) with the R Studio IDE v1.4.1717 (www.rstudio.com/). Sequencing reads were demultiplexed, filtered, and trimmed using Cutadapt v.4.1 (Martin, 2011); adapter sequences (e.g., AGATCGG), as well as poly-A and poly-G tails that exceeded 8 base pairs (bp) were removed. The first 15 bp of the 3’ end of each fragment was trimmed to prevent the inclusion of low-quality reads (quality-cutoff). After trimming, any fragment that was less than 20 base pairs were removed (minimum read length). Trimmed reads were aligned to the Roslin Institute’s *Crassostrea gigas* genome assembly with mitochondrial genes added (Peñaloza et al., 2021) using Hisat2 v.2.2.1 (D. Kim et al., 2019), a splice aware aligner that has previously been used effectively with Tag-Seq data generated from bivalves (Gurr et al., 2022). Aligned reads were assembled against the mRNA genome feature track of the Roslin genome, merged using StringTie v2.2.0, and compiled using prepDE.py (Kovaka et al., 2019). Gene count matrices were analyzed with DESeq2 v.1.36.0 (Love et al., 2014) using the apeglm shrinkage estimator (Zhu et al., 2019). Genes with counts of more than 10 across at least one-third of samples were included in downstream analyses. Significant DEG lists were generated for each comparison using a Wald test p-value cutoff of 0.05.

### 2.8. Functional Enrichment Analysis

Functional enrichment analysis was performed to identify gene ontology (GO) terms significantly overrepresented within ploidy-specific DEG lists. Gene lists were analyzed for diploids and triploids within the single and multiple stressor treatments. To obtain GO terms associated with each DEG, uniprot Accession information for RNA nucleotide sequences (transcript IDs) was first obtained through a BLAST (Altschul et al., 1990) of the *C. gigas* genome (Peñaloza et al., 2021) against the Uniprot-SwissProt database (UniProt Consortium, 2019). A gene ID-to-GO term database was created by matching GO terms from the Uniprot-SwissProt database to transcript IDs from the blast output using Uniprot Accession codes; GO terms associated with each DEG transcript ID were identified using this database.

Gene enrichment analysis was conducted using GOseq v.1.48.0 in R (Young et al., 2010). The Wallenius approximation was used to identify over-represented GO terms within the Biological Processes gene ontology branch, using all measured genes from individuals included in each analysis as a background. Enriched (over represented) GO terms were filtered using an adjusted p-value cutoff of 0.05, de-duplicated, and assigned a functional category. Functional category assignment was performed by categorizing identified GO terms into subgraphs (GOcats) based on a curated list of user-generated terms of interest (Hinderer & Moseley, 2020); Selected terms included cellular processes (including response to stimulus, transport), regulation (biological, epigenetic), metabolism (protein, RNA, other), immune system processes, and stress response (including homeostasis, death). GOterms that failed to match to any defined GOcat were assigned to other biological processes. The distribution of enriched GOterms within each functional category were compared with a chi-squared test, using the results obtained for diploids within the single or multiple stressor treatment as the expected values.

### 2.9. Statistical analyses

All statistical analyses were performed in R v.4.2.1 using the RStudio IDE v.2022.7.1.554. When applicable, analysis of variance (ANOVA) was used to investigate differences in response variables (e.g., ATPase, citrate synthase) across stress treatments. For response variables with multiple observations from the same individual (e.g., metabolic rate), a repeated measures within-subject analysis of covariance (ANCOVA) was used to compare the impact of each factor (ploidy, stress treatment, duration of exposure), using oyster ID as a random effect. Assumptions of normality and homoscedasticity were assessed during model construction using the Shapiro test and a visual assessment of Q-Q and residual-fitted plots. The Johnson transformation was used to achieve normality when necessary (Fernandez & Fernandez, 2012). For significant effects or their interactions (α = 0.05), the agricolae package was used to perform pairwise comparisons of groups using the Tukey HSD post hoc test (Mendiburu & Yaseen, 2020).

## 3. Results

Ploidy analysis revealed that 100% of oysters tested (n=30) corresponded to their presumed ploidy. The shell length, height, and width of oysters significantly varied across ploidy (p<0.001), but not across treatment (Table S1–S2). Despite size differences, the dry tissue weight of oysters did not vary across ploidy or treatment (Table S2). Upon arrival at the hatchery, reproductive condition significantly varied across ploidy, with a larger proportion of diploids exhibiting later stage gonad development than triploids (*X*^2^_df = 1, N = 24_ = 78.1, p<0.001); this difference was maintained after single stress exposure (*X*^2^_df = 1, N = 25_ = 352, p<0.001). However, no difference across ploidy was observed following multiple stress exposure (*X*^2^_df = 1, N = 25_ = 9.57, p=0.08). At day 30 within the control treatment, there was also no significant difference in reproductive condition across ploidy (*X*^2^_df = 1, N = 24_ = 1.83, p=0.87). The proportion of diploid and triploid oysters at each reproductive stage at each time point is presented in Figure 1D.

### 3.1. Treatment conditions

Seawater temperature significantly varied across treatment (p<0.001), with the single and multiple stressor treatments maintaining an elevated temperature with respect to the control for thirty days following the acclimation period (Figure S4). The pH and salinity were not significantly different across treatments. The average (± SD) for each measured seawater parameter is presented in Table 1.

Diploid and triploid oysters within the multiple stressor treatment group were subjected to aerial emersion within one of four custom-built desiccation chambers on day 2 for 6 h. Oysters were afforded a 1 h ramp and cooling period preceding and following the maintenance of each chamber at a temperature setpoint of 44°C (Figure 1B). The mean air temperature within the maintenance period across chambers was 44.9 ± 1.6°C, corresponding to a mean relative humidity of 34.0 ± 4.0%. The mean internal temperature of oysters distributed across desiccation chambers was 33.9 ± 4.0°C while temperature was being maintained (n=7; Figure 1B). The maximum internal temperature of oysters significantly varied with shell volume (p=0.04, r^2^=0.59, Figure 1C). The highest internal temperature recorded across all oysters monitored was 47.3°C.

### 3.2. Survival Analysis

Mortality was observed within all treatment groups over the course of the experiment. The mortality rate for both diploids and triploids were elevated within the stress treatments with respect to the control and was significantly higher in triploids than in diploids following multiple stress exposure (Figure 2A-B). The cumulative mortality across treatments at day 30 was 5.7%, 9.1%, 14.8%, 22.7%, 9.1%, and 36.4% across the 2n-C, 2n-SS, 2n-MS, 3n-C, 3n-SS, and 3n-MS treatments, respectively. Survival probability was significantly impacted by ploidy (p = 0.017) and stress treatment (p<0.001). Triploids within the multiple stressor treatment (3n-MS) had the lowest 1-month survival probability (0.7 ± 0.09), followed by the 2n-MS (0.86 ± 0.07), 3n-SS (0.88 ± 0.07), 3n-SS (0.93 ± 0.05), 3n-C (0.94 ± 0.05), and 2n-C (0.95 ± 0.05) treatments, respectively.

**Figure 2.**
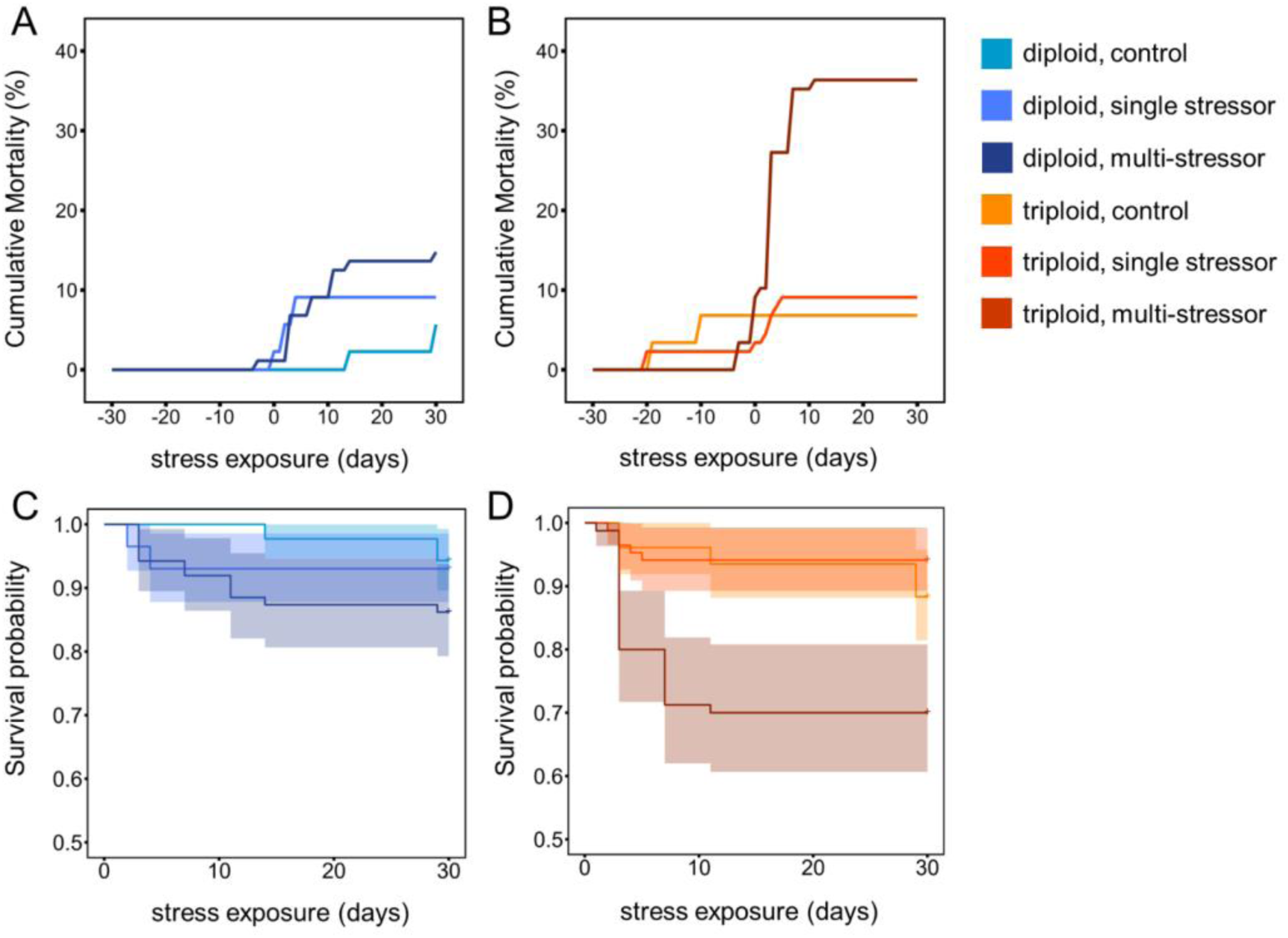
Cumulative mortality (A-B) and survival probability (C-D) for diploid and triploid adult Pacific oysters within hatchery stressor trials. All oysters were acclimated to hatchery conditions (day −30 to −10, 20°C) before inclusion in either the control (20°C), single stressor (SW temp = 30°C), or multiple stressor (SW temp = 30°C; aerial exposure = 44°C for 4 h on d2) treatment.

### 3.3. Standard Metabolic Rate

The metabolic rate of oysters was significantly affected by the interaction of ploidy (diploid, triploid), stress treatment (control, single stressor, multiple stressor), and the duration of stress exposure (p<0.001; Table S3). Metabolic rate did not vary significantly across ploidy within the control treatment (20°C), and was similar during the acclimation period (day −30 to −10; “before experiment) and on day 10 (“after experiment”; Figure 3A). For diploids, metabolic rate was significantly elevated following multiple stressor exposure with respect to the single stressor treatment; this difference was maintained for up to 8 days after multiple stress exposure (Figure 3B,D). In contrast, no significant increase in metabolic rate was observed in triploids following multiple stressor exposure (Figure 3C,E). When compared with diploids, triploids experienced metabolic depression following MS exposure (Figure 3F).

**Figure 3.**
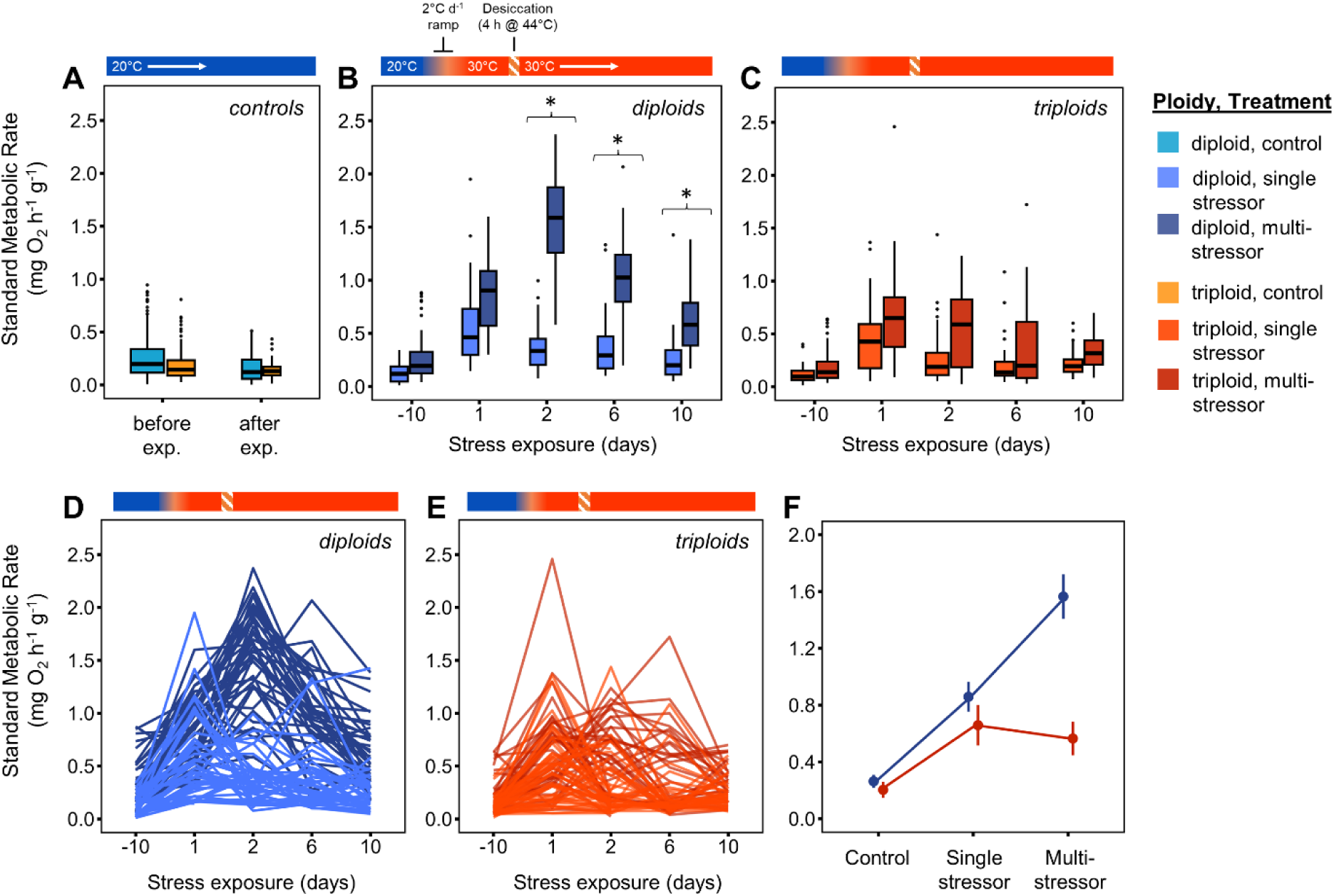
The impact of ploidy, stress type, and exposure duration on the standard metabolic rate of adult Pacific oysters. Oysters were acclimated to hatchery conditions (day −30 to −10, 20°C) before inclusion in either the control (20°C), single stressor (SW temp = 30°C), or multi-stressor (SW temp = 30°C; aerial exposure = 44°C for 4h on d2) treatment. Italicized letters represent the result of post-hoc Tukey HSD comparisons across groups; boxplots that share a letter within panels A-C are not significantly different. Oxygen consumption was repeatedly sampled from individually labeled oysters (n=30-45 per treatment), each of which are represented as lines in panels D and E. The mean metabolic rate (± 95% CI) of oysters preceding and directly following multiple stress exposure are presented in panel F.

### 3.4. Enzymatic assays

Na^+^/K^+^ ATPase activity was significantly impacted by the interaction of ploidy (diploid, triploid), stress treatment (control, single stressor, multiple stressor), and the duration of stress exposure (p=0.009; Table S4). ATPase activity was significantly lower in triploids than diploids after multiple stressor exposure (Figure 4A; Figure S5). Citrate synthase was significantly impacted by stress treatment (p<0.001; Table S4), with greater activity observed in the multiple stress treatment than the single stressor and control within both diploids and triploids (Figure 4B). Citrate synthase activity was not significantly impacted by ploidy (p=0.487; Table S4).

**Figure 4.**
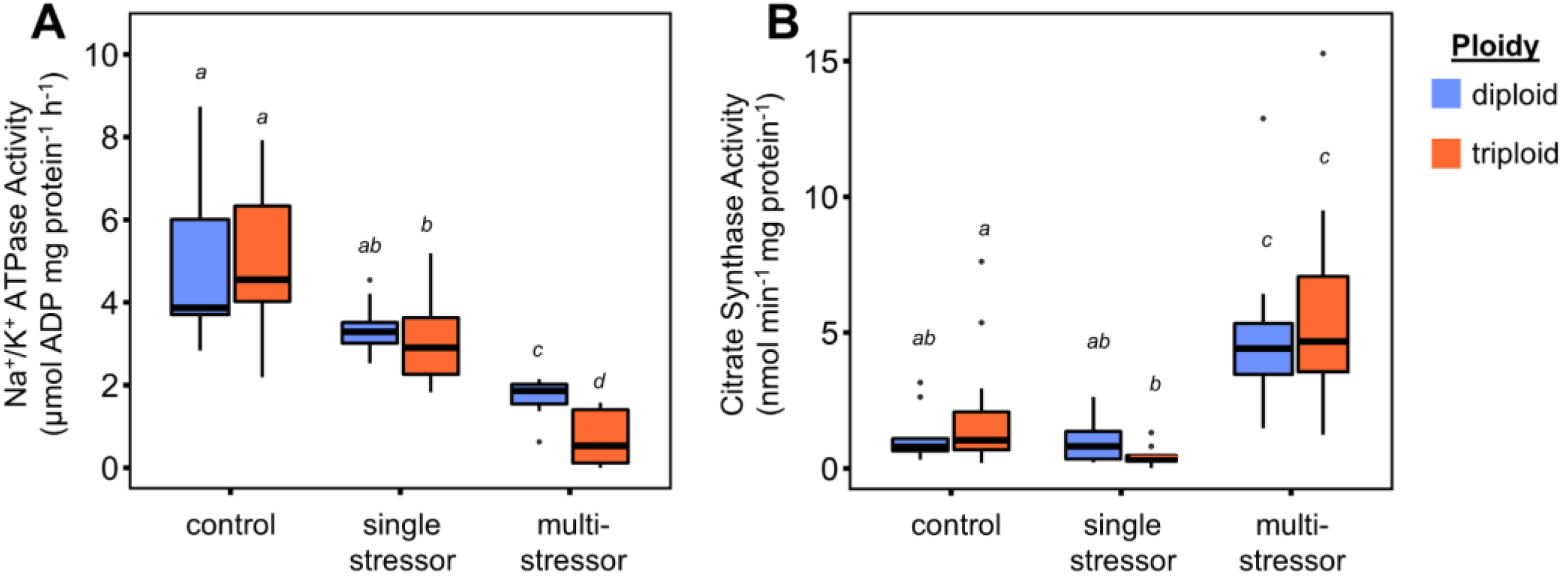
The impact of ploidy and stress exposure on Na^+^/K^+^ ATPase (A; n=10-13 per treatment; 1d after stress) and citrate synthase (B; n=9-12 per treatment; 1d after stress) enzyme activity of adult Pacific oysters. All oysters were acclimated to hatchery conditions (20d, 20°C) before inclusion in either the control (20°C), single stressor (SW temp = 30°C), or multi-stressor (SW temp = 30°C; aerial exposure = 44°C for 4h) treatment. Italicized letters represent the result of post-hoc Tukey HSD comparisons across groups; boxplots that share a letter within panels are not significantly different. All enzyme activity was measured within the ctenidium.

### 3.5. Differential gene expression (DEG) analysis

Out of the seventy-two biological samples submitted for sequencing, eight were removed prior to analysis for quality concerns following the inspection of FastQC reports (Ewels et al., 2016). Sample sizes after quality control were 11, 11, 11, 10, 10, and 11 for the 2n-C, 3n-C, 2n-SS, 3n-SS, 2n-MS, and 3n-MS treatment, respectively. Remaining samples averaged 4.94 ± 1.1 million reads and achieved an average alignment rate of 87.8 ± 1.3% with *Crassostrea gigas* genome (GenBank: GCA_902806645.1).

The *DESeq* function identified 4258, 1300, 594, and 668 significant DEGs associated with the 2n-SS, 3n-SS, 2n-MS, and 3n-MS treatment, using the control treatment within each corresponding ploidy as the baseline condition; full identified gene lists for each comparison are provided in Table S5-S8. Significant expression differences were observed across a suite of genes that encode molecular chaperones (Table 2) and proteins that function to inhibit or regulate cellular apoptosis (Table 3).

**Table 2.** Differentially expressed genes (DEG) encoding molecular chaperones within diploids and triploids following single or multiple stress exposure.

| GeneID | Description | Log2FoldChange |  |  |  |
| --- | --- | --- | --- | --- | --- |
|  |  | 2n-SS | 3n-SS | 2n-MS | 3n-MS |
| 105345220 | activator of 90 kDa heat shock protein ATPase | 1.1 ± 0.3 |  |  |  |
| 105334580 | BAG family molecular chaperone regulator 3 |  |  | 5.1 ± 0.7 | 3.4 ± 0.5 |
| 105328795 | heat shock protein Hsp-16.2 |  |  | 1.5 ± 0.5 |  |
| 105343216 | heat shock protein 27 | 3.1 ± 1 |  | 5.4 ± 0.5 | 3.4 ± 0.5 |
| 105348623 | heat shock protein 60A | 0.6 ± 0.2 |  |  |  |
| 105334510;<br>105348304 | heat shock protein 68 | -0.9 ± 0.4;<br>- |  | 8.8 ± 0.7;<br>8.9 ± 0.7 | 8.6 ± 0.8;<br>8.2 ± 0.8 |
| 105322484;<br>105348305;<br>117680673 | heat shock protein 68-like |  |  | 5.8 ± 0.6;<br>8.4 ± 0.7;<br>8.3 ± 0.7 | 5.7 ± 0.6;<br>8.0 ± 0.8;<br>8.2 ± 0.8 |
| 105344160;<br>105349016 | heat shock protein 70 kDa 12A | 0.6 ± 0.3<br>-1.4 ± 0.7 |  |  |  |
| 117682412;<br>105324023;<br>117681885 | heat shock protein 70 kDa 12A-like | 0.6 ± 0.2;<br>-1.2 ± 0.4<br>-1.1 ± 0.3 | 0.8 ± 0.4;<br>-;<br>- | 0.6 ± 0.2;<br>-;<br>- |  |
| 105334234 | heat shock protein 70 B2 | 0.9 ± 0.7 |  | 5.9 ± 0.7 | 3.9 ± 0.6 |
| 105326385 | heat shock 70 kDa protein 4 | 0.6 ± 0.2 | 0.6 ± 0.3 |  |  |
| 105342539 | heat shock 70 kDa protein 14 | 0.6 ± 0.2 |  |  |  |
| 105345989;<br>117680291 | heat shock protein 83-like | 2 ± 0.2;<br>1.9 ± 0.2 | 2 ± 0.3;<br>1.9 ± 0.3 | 1.1 ± 0.3;<br>1.1 ± 0.3 |  |
| 105321527 | peptidyl-prolyl cis-trans isomerase B | 0.6 ± 0.2 |  |  | 0.4 ± 0.2 |
| 105321016 | protein canopy homolog 3 | 1.1 ± 0.2 | 0.9 ± 0.1 | 1.2 ± 0.2 | 0.7 ± 0.1 |
| 105346349 | protein disulfide-isomerase A3 | 1.2 ± 0.2 | 0.8 ± 0.2 | 0.8 ± 0.2 |  |
| 105322320 | protein disulfide-isomerase A4 isoform X2 | 1.6 ± 0.2 | 1.3 ± 0.2 | 1.2 ± 0.2 | 0.9 ± 0.2 |
| 105341388 | protein disulfide-isomerase A6 isoform X1 | 1.6 ± 0.2 | 2 ± 0.2 | 0.7 ± 0.3 | 0.9 ± 0.2 |
| 105330729 | protein SGT1 homolog | 0.7 ± 0.2 | 0.5 ± 0.2 |  | 0.4 ± 0.2 |

**Table 3.**
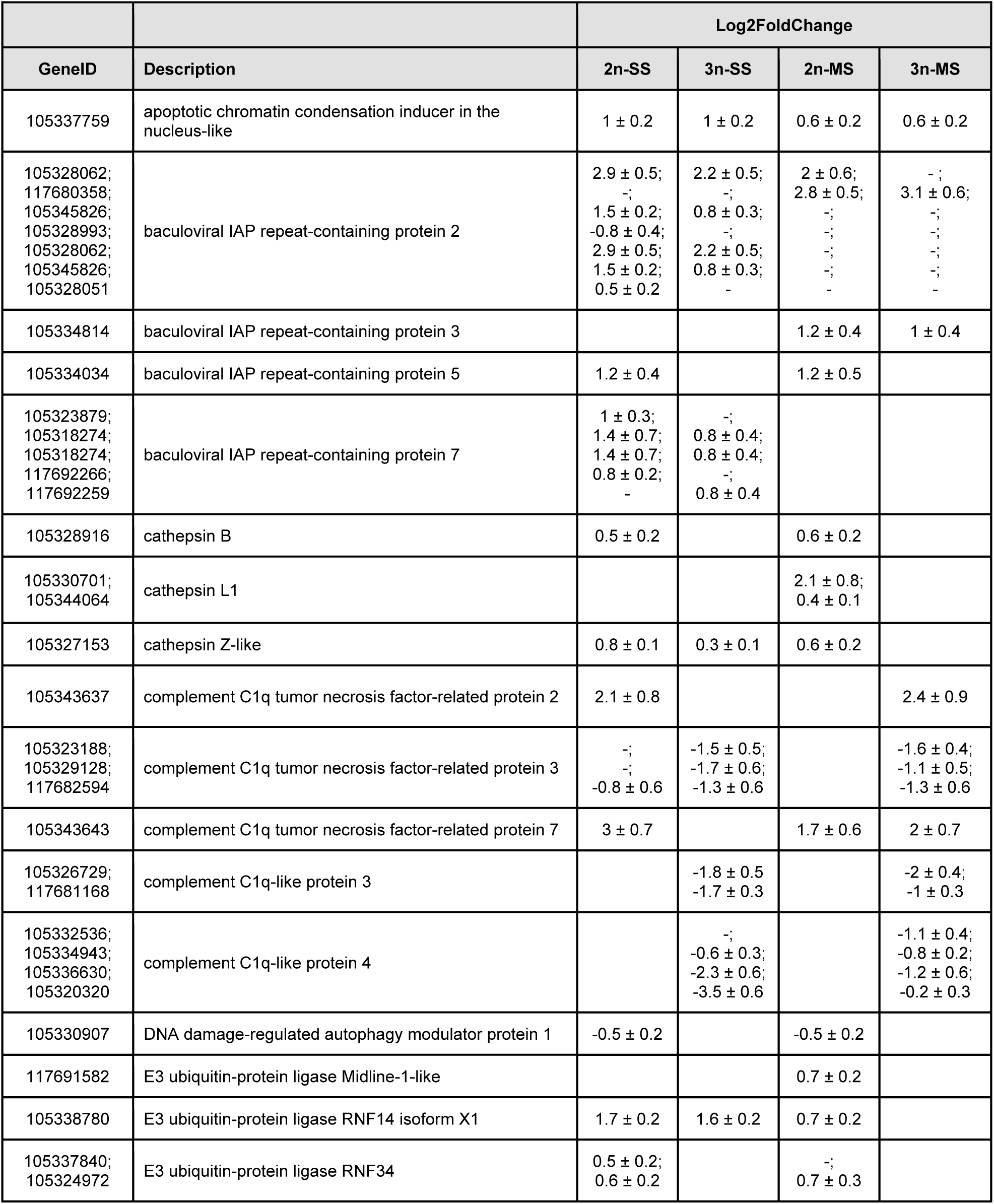
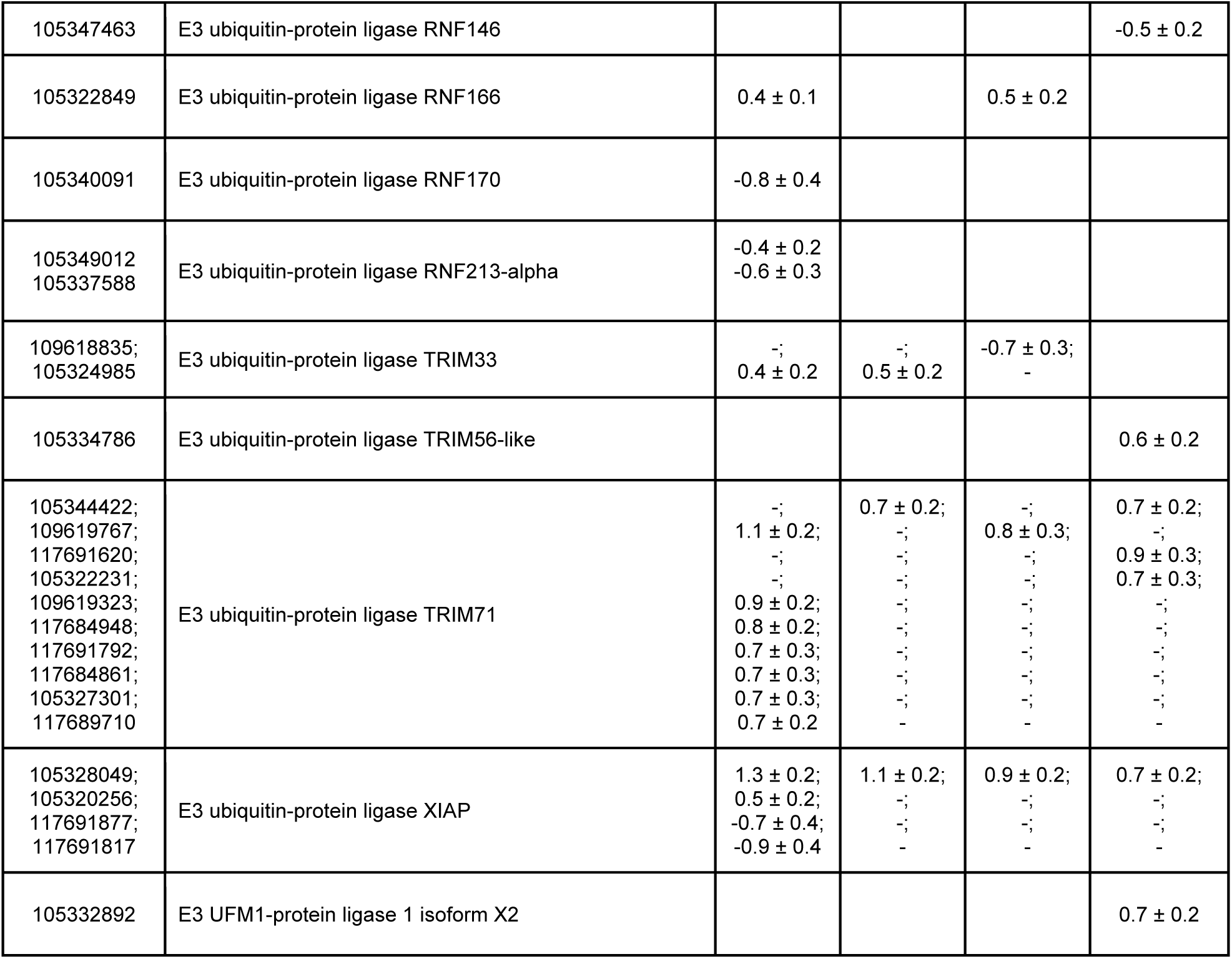
Differentially expressed genes (DEG) whose protein products function to inhibit or regulate cellular apoptosis within diploid (2n) and triploid (3n) following single (SS) or multiple (MS) stress exposure.

Principal component analysis demonstrated that the gene expression profiles of diploids within the single and multiple stressor treatments were more similar to each other than to the control (Figure 5B). In contrast, the gene expression profiles of triploids within the single and multiple stressor treatments overlapped considerably with the control (Figure 5C). Across ploidy, triploids exhibited a greater degree of gene expression variation than diploids prior to being stressed (Figure 5D). The gene expression profiles of diploids and triploids varied considerably following stress exposure, with marginal overlap between individuals following exposure to elevated seawater temperature (Figure 5E) and no overlap following exposure to elevated seawater temperature and desiccation in combination (Figure 5F).

**Figure 5.**
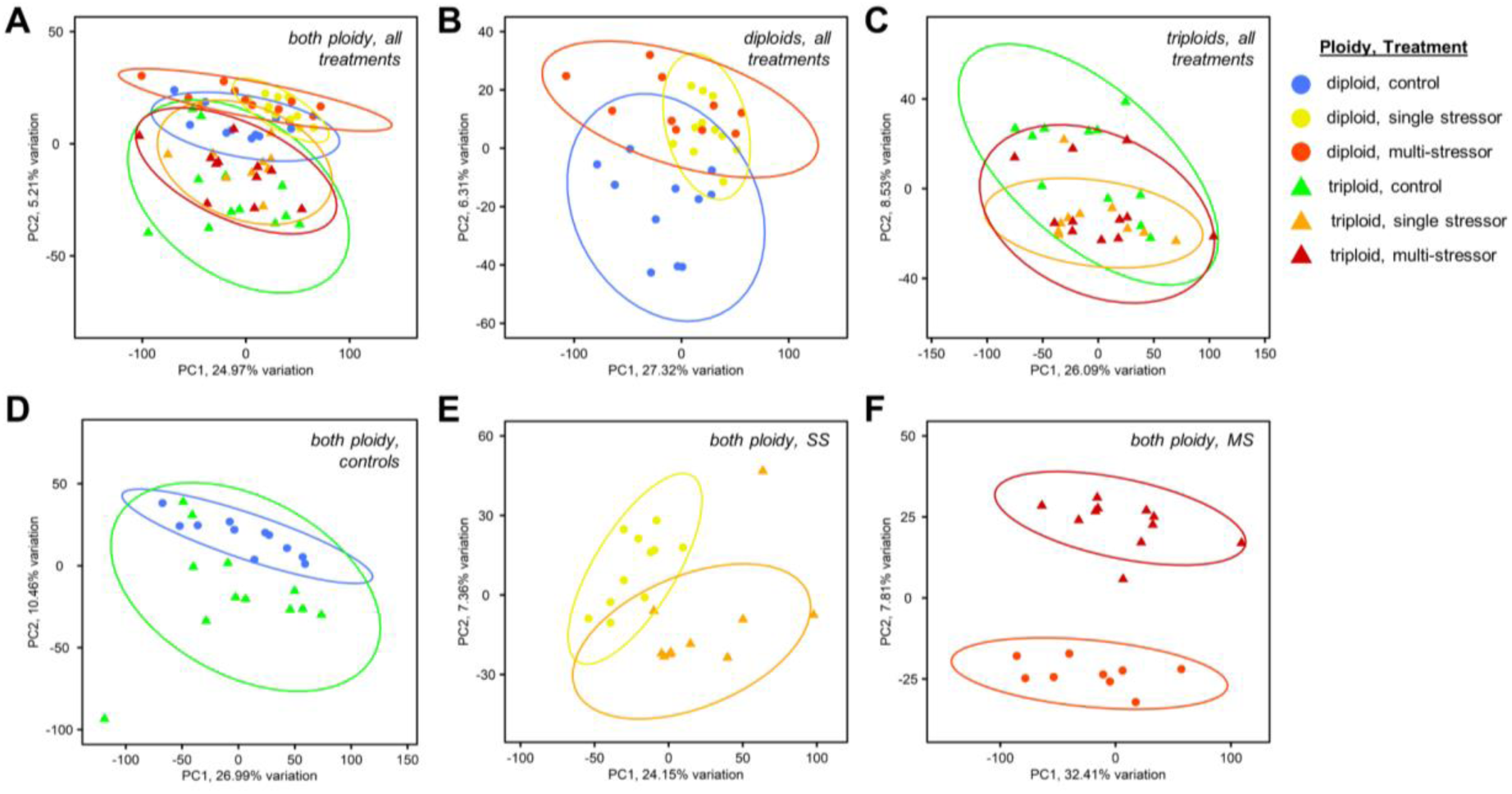
Principal component plots comparing gene expression across all treatments (A), within ploidy (B, C), or within the control (D), single stressor (E), or multi-stressor (F) treatment.

The number of unique and overlapping DEG associated with environmental stress within diploid and triploid oysters varied across treatments. A 6-fold greater number of unique DEG were observed in the 2n-SS treatment (3368) than the 3n-SS treatment (548), while a comparable number was observed in the multiple stressor treatment (2n: 159; 3n: 210; Figure 6A). Under control conditions, a nearly equal proportion of DEG exhibited either increased or decreased expression across ploidy (Figure 6B). Following multiple stress exposure, this trend was maintained in triploids (Figure 6E), while diploids exhibited a greater number of genes with decreased expression (Figure 6D). A cluster analysis across each comparison confirmed that DEG expression patterns were more similar within, than across, ploidy (Figure 6A) and within treatments (Figure 6D,E).

**Figure 6.**
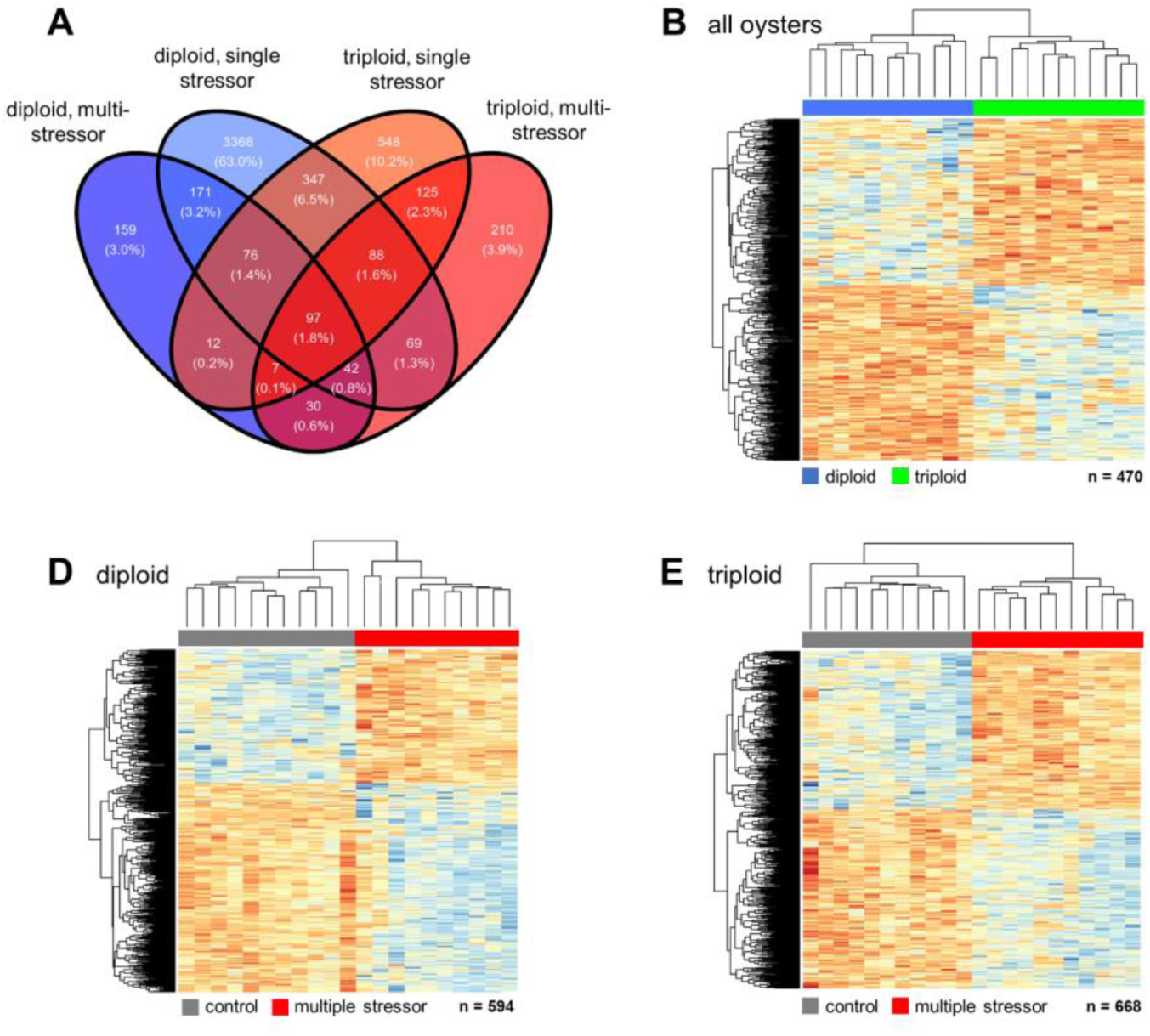
(A) Venn diagram of overlapping and contrasting differentially expressed genes (DEG) identified within comparisons of treatment (SS: single stressor; MS: multi-stressor) and ploidy (diploid; triploid). Z-score heatmaps of significant DEG within the controls across ploidy (C) and between the control and multi-stressor treatment for diploids (D) and triploids (E).

### 3.6. Functional Enrichment Analysis

Ploidy-specific DEG from the single stressor and multiple stressor treatments were used in functional enrichment analysis. Following single stressor exposure, 3650 DEG were uniquely associated with diploids, while 692 were associated with triploids (Figure 7A). Significant single stressor DEG mapped to 100 diploid-specific and 66 triploid-specific enriched GOterms associated with biological processes (Figure 7B); the distribution of GOterms within each functional category was significantly different across ploidy (*X*^2^_df = 6, N = 65_ = 37.41, p<0.001) following single stressor exposure, with 11.9% and 29.2% of terms associated with stress and immune processes within diploids and triploids, respectively. Following multiple stressor exposure, 418 DEG were uniquely associated with diploids and 492 were associated with triploids (Figure 7C). Significant multiple stressor DEG mapped to 74 diploid-specific and 50 triploid-specific enriched GOterms associated with biological processes (Figure 7D); the distribution of GOterms associated with functional categories was also significantly different across ploidy (*X*^2^_df = 6, N = 50_ = 38.79, p<0.001), with 10.81% of diploid-specific and 26.0% of triploid-specific terms associated with stress and immune processes.

**Figure 7.**
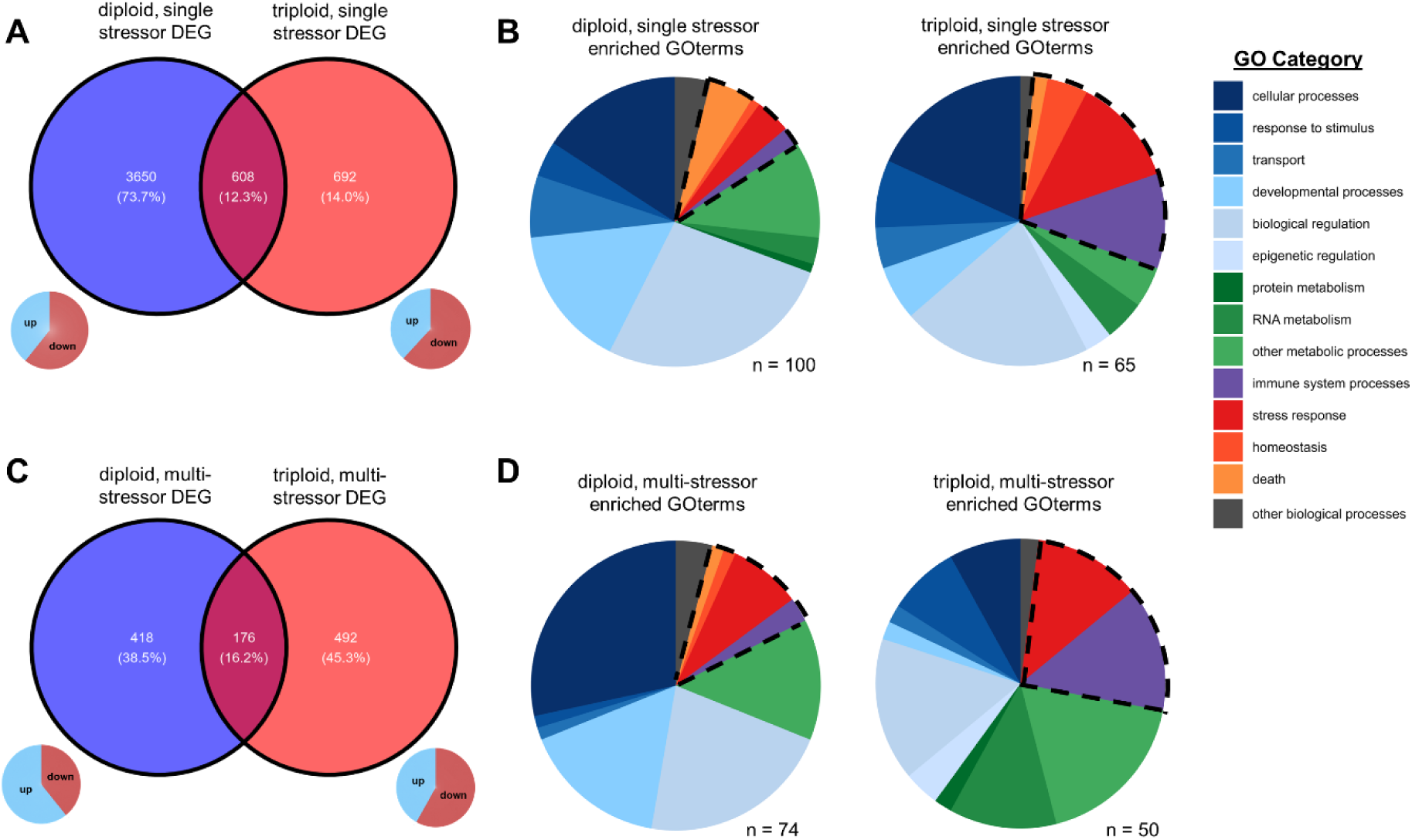
Venn diagram of overlapping and contrasting significant differentially expressed genes (DEG) across ploidy within the single stressor (A) and multi-stressor (C) treatment. (B) Pie charts of significantly enriched GOterm categories associated with unique DEG within each ploidy after exposure to single (B) and multiple stressors (D).

## 4. Discussion

The primary objective of this study was to determine the impact that polyploidy has on environmental stress tolerance, using the response of Pacific oysters to marine heatwaves (MHW) as a model system. In this study, the mortality rate of diploid and triploid oysters was similar after a gradual elevation in seawater temperature to 30°C (single stressor). However, when followed by aerial emersion that replicated conditions present during a MHW (multiple stressors, 44°C aerial temperature), triploids exhibited a 2.5-fold higher mortality rate than diploids (36.4% vs. 14.8%). An elevated mortality in triploids was paired with metabolic rate depression, the depletion of Na+/K+ ATPase within the ctenidium, and the dysregulated expression of genes the encode heat-responsive molecular chaperones, antiapoptotic proteins, and mitochondrial genes associated with aerobic metabolism. These results suggest that polyploidy alters the genetic architecture that regulates the stress response in *C. gigas*, negatively impacting physiological tolerance limits and increasing susceptibility to climate events like MHWs.

### 4.1. Triploid oysters display higher mortality rates within multiple stressors scenarios

While the thermotolerance of *C. gigas* is well studied, the impact of polyploidy on thermal tolerance limits remains unclear. Thermal limits in oysters vary across populations and can be plastic, with recent work demonstrating shifts in tolerance can occur following prior exposure to marine heatwaves (Ding et al., 2020; He et al., 2021). Despite this variability, diploid Pacific oysters generally switch between aerobic and anerobic metabolism between 36-43°C when submerged, with 100% mortality observed after this range is exceeded (Ghaffari et al., 2019). A small, but growing, number of laboratory experiments suggest that triploids have a lower critical thermal maximum than diploids; For example, (Li et al., 2022) found that triploid *C. gigas* display a 1.51-fold higher cumulative mortality rate than diploids (57.05% vs. 37.73%) following acute heat shock (18°C to 28°C) and a 3.25-fold higher mortality rate (83.7% vs. 25.73%) following chronic heat exposure. However, field assessments of triploid and diploid oyster survivorship over the course of a growing cycle suggest that summer mortality is more complicated than would be expected from differences in thermal tolerance limits alone. Triploid *C. gigas*, *C. virginica*, and *C. hongkongensis* have been shown to exhibit lower (Gagnaire et al., 2006; Qin et al., 2019), higher (Bodenstein et al., 2021, 2023; Guévélou et al., 2019; Houssin et al., 2019; Matt, 2018; Matt et al., 2020; Wadsworth, 2018; Wadsworth et al., 2019), and comparable (Dégremont et al., 2012; Ibarra et al., 2017) mortality rates when compared with diploids depending on the environmental conditions present. For example, Wadsworth et al. (2019) found higher cumulative mortality in triploid *C. virginica* than diploids across growing sites within the Gulf of Mexico; mortalities were associated with sudden drops in salinity (< 5 ppt) and high temperature (> 30°C). The presence of multiple stressors preceding mortality events is consistent with both Guévélou et al. (2019) and Matt et al. (2020), who have noted that *C. virginica* triploid mortalities observed within Chesapeake Bay, VA are not correlated with the presence of elevated seawater temperature alone. Instead, common farm stressors, including tumbling stress and desiccation, as well as hyposalinity have been shown to contribute to triploid mortality (Bodenstein, 2019; Bodenstein et al., 2021, 2023). These results are consistent with the findings of this study that suggest that the synergistic impact of multiple environmental stressors can result in ploidy-specific differences in energy balance that contribute to differential survival.

### 4.2. Summer mortality as a reproductive disorder

While energy allocation to gametogenesis may play a role in the initiation of summer mortality events under field conditions, there was no evidence of an interaction between reproductive investment and mortality within this study. To replicate the seasonal timing of marine heatwaves, hatchery experiments took place in summer (June-July) and were intentionally timed to coincide with the period of peak reproductive output. While large-scale summer die-offs within oysters are common and have been associated with a variety of abiotic and biotic factors, summer mortality has also been interpreted as a reproductive disorder (Koganezawa, 1975; Mori, 1979). Shifts in carbohydrate storage and utilization leading up to spawning can place energetic constraints on metabolism; as such, reproductive effort has been associated with higher mortality rates during the spring and summer in diploids, especially under elevated temperature and in the presence of marine pathogens (Huvet et al., 2010; Wendling & Wegner, 2013). Histological examination of the reproductive tissue of triploid oysters in this study confirmed early (stage 1) or no gonad development (stage 0) throughout the experiment; in comparison, the majority of diploids had mature gonads (stage 3) upon arrival at the laboratory and after single stress exposure, with evidence of spawning only after multiple stress exposure (Figure 1D). Given that the mortality rate of triploids exceeded that of diploids following multiple stress exposure, it is unlikely that reproductive state was the motivating factor that impacted stress tolerance. However, it should be noted that although triploid oysters are commonly referred to as sterile, they can produce gametes (Allen & Downing, 1990; Jouaux et al., 2010; Matt & Allen, 2021), a phenomenon that may be enhanced by warm seawater temperatures (Normand et al., 2008). Future studies would benefit from an in depth investigation of whether the incomplete formation or rupture of gonads during gametogenesis contributes to increased rates of bacterial infection and mortality within triploids (De Decker et al., 2011).

### 4.3. Triploid mortality as a physiological syndrome

The ‘MOREST’, a multi-disciplinary program coordinated by IFREMER to investigate oyster mortalities within France, concluded that summer mortality may be a ‘physiological syndrome’ that results from energetic limitation under multiple stressor scenarios (Samain & McCombie, 2008). While this may be true for diploids as well, the results of multiple physiological assays employed in this study suggest that triploids experience a greater degree of energetic limitation than diploids following exposure to multiple stressors. First, triploids, but not diploids, exhibited metabolic depression that persisted for up to 10 days following multiple stress exposure (Figure 3). Metabolic depression following an environmental stress can be protective over short durations by effectively reducing ATP and protein turnover rates to maintain energy balance. However, a sustained reduction in oxygen consumption while under physiological stress can result in the accumulation of anaerobic metabolites and the discontinuation of important ATP-demanding processes that can, in turn, increase mortality rates (Guppy & Withers, 1999; Lesser, 2016; Zittier et al., 2015). Second, triploids experienced a significantly greater depletion of ctenidium Na^+^/K^+^ ATPase enzyme activity following multiple stressor exposure than diploids (Figure 4A). Na^+^/K^+^ ATPase is a ubiquitously expressed multifunctional transmembrane protein complex that is essential for numerous physiological processes such as osmotic and ionic regulation, calcification, and ammonia excretion in animals (Crane, 1977; Skou, 1957; Wright & Manahan, 1989) and is commonly used an indicator of energetic limitation following environmental stress exposure (Bianchini & Wood, 2003; Wheatly & Henry, 1987). Na^+^/K^+^ ATPase mediated physiological processes are energy intensive, accounting for 20% of the energy expenditure in mammalian tissues (Milligan & McBride, 1985) and up to 77% of larval oxygen consumption in the sea urchin *Strongylocentrotus purpuratus* (Leong & Manahan, 1997).

A precipitous decrease in Na^+^/K^+^ ATPase enzyme activity following multiple stress exposure is consistent with triploids exhibiting metabolic rate depression and transitioning into ‘pessimum’ or a state of time-limited survival (Sokolova et al., 2012). Within the pessimum range, all available metabolic energy is reallocated to somatic maintenance; using this mechanism, metabolic demand is reduced in an attempt to ‘wait out’ an environmental stressor. Given the enhanced carbohydrate reserves of triploids (Shpigel et al., 1992), it is surprising from a bioenergetics perspective that triploids, but not diploids, display metabolic arrest and a higher mortality rate following multiple stress exposure. One possible explanation is that triploidy negatively impacts the morphology or function of the heart. Several triploid salmonids have been shown to experience cardiac arrhythmias at lower temperatures than diploids, resulting in inadequate oxygen transport, a reduction in factorial metabolic scope, and a reduction in thermal limits (Altimiras et al., 2002; Fraser et al., 2013; Verhille et al., 2013). The cardiac response of bivalves varies with thermal variability and thermal history, making it a useful metric in the determination of metabolic capacity (Nancollas & Todgham, 2022). Future studies would benefit from an in-depth investigation into the cardiac responses of oysters preceding and following marine heatwaves to ascertain whether the reduction in metabolic rate in triploids observed in this study corresponds to ploidy-level differences in the onset of cardiac failure following exposure to multiple environmental stressors.

### 4.4. Triploids exhibit signs of transcriptional dysregulation following multiple stress exposure

One explanation that could account for ‘triploid mortality’ is the dysregulation of key biological processes due to changes in gene dosage, the genetic architecture of regulatory networks, incomplete silencing, epigenetic instability, and/or other downstream impacts of chromosome set duplication (Comai, 2005). Dysregulated gene expression can negatively impact the function and efficiency of gene networks, resulting in energetic costs and disease (Y.-A. Kim et al., 2011). However, if the energetic cost of a gene’s dysregulation is minor, any negative consequences or downstream impacts of energy budgets may only manifest when multiple stressors are present. For example, *C. virginica* is remarkably tolerant to a range of environmental stressors, but the simultaneous exposure of two or more stressors can reduce the tolerance to any single factor (Cherkasov et al., 2006; Ivanina et al., 2012; Kurochkin et al., 2009). The effects of two or more stressors are considered synergistic if an organism’s response to them together is greater than the sum of their responses independently (Todgham & Stillman, 2013). Synergistic effects following multiple stressor exposure have been reported in several oyster species, including interactions between warming and salinity (Jones et al., 2019; Marshall et al., 2021; McFarland et al., 2022) and hypoxia and ocean acidification (Gobler et al., 2014).

Functional enrichment analysis of ploidy-specific gene sets indicated that GO terms associated with stress and immune processes were significantly enriched in triploids following single and multiple stress exposure (Figure 7). This result is consistent with the findings of Li et al. (2022), who found an overrepresentation of inflammatory response and apoptosis gene expression following acute temperature stress in triploid *C. gigas*. In this study, triploids exhibited dysregulated expression of stress-related proteins following multiple stress exposure, with notable differences in three classes of proteins: molecular chaperones and heat shock proteins, inhibitor of apoptosis (IAP) proteins, and E3 ubiquitin-protein ligases. HSPs are a broad class of molecular chaperones that are capable of mediating heat-induced cellular damage by preventing protein misfolding (Meistertzheim et al., 2007). Following multiple stress exposure, diploids increased the expression of eleven genes associated with HSPs, while triploids only expressed seven, the list of which omitted HSP-16.2 (GeneID: 105328795) and HSP-83 (GeneID: 105345989, 117680291; see Table 2). IAPs bind to and block tumor necrosis factor receptor–associated factors (TRAFs) to prevent cell death (Rothe et al., 1995). The expression patterns of a variety of IAPs varied across ploidy, with notable differences in the expression of baculoviral IAP repeat-containing protein −2 (GeneID: 105328062) and −5 (GeneID: 105334034), as well as complement C1q tumor necrosis factor-related protein 3 (GeneID: 105323188; 105329128; 117682594), complement C1q-like protein 3 (GeneID: 105326729; 117681168) and −4 (GeneID: 105332536; 105334943; 105336630; 105320320; see Table 3). Similarly, elevated expression of a suite of E3 ubiquitin-protein ligases species was observed in diploids, but not in triploids, including E3 ubiquitin-protein ligase Midline-1 (117691582), RNF14 (105338780), RNF34 (105324972), RNF166 (105322849), and TRIM33 (109618835). E3 ubiquitin-protein ligases are known to protect against apoptosis by regulating p53 protein, a protein that plays a pivotal role in the initiation of cell division and cell death (M. Pan & Blattner, 2021).

### 4.5. Triploids exhibit key differences in the expression and activity of mitochondrial enzymes involved in aerobic metabolism following multiple stress exposure

In addition to stress-response proteins, significant differences in the expression patterns of a variety of genes that regulate mitochondrial function and aerobic metabolism were observed across ploidy following multiple stress exposure. Elevated expression of methyltransferases and other proteins involved in stress-related mitochondrial biogenesis and proliferation was observed in diploids, but not triploid, such as rRNA methyltransferase 2 (GeneID: 105338531, Rorbach et al., 2014), and mitochondrial genes implicated in heat and oxidative stress tolerance, including NADH-ubiquinone oxidoreductase (GeneID: 105342980, Downs & Heckathorn, 1998), 39S ribosomal protein L19 (GeneID: 105320404, Chen et al., 2020) and isocitrate dehydrogenase [NADP] (GeneID: 105343391, Jo et al., 2001). The elevated expression of genes with pivotal regulatory roles in the TCA cycle were also only observed in diploids, including cytochrome P450 (GeneID: 105336572, 105338167), cytochrome b-c1 (GeneID: 105345808, 105317345), and cytochrome c oxidase (GeneID: 105340053). The muted transcriptional response of triploid oysters observed in this study is in agreement with prior work that suggests that triploid *C. gigas* are less sensitive to environmental cues than diploids (Duchemin et al., 2007) and mount a delayed gene regulatory response to heat shock within laboratory assays (Li et al., 2022). Additionally, evidence of dysregulated expression of nuclear genes that dictate mitochondrial activity and energy production in polyploids supports the hypothesis that triploids are forced into a state of energetic limitation within multiple stressor scenarios.

Interestingly, citrate synthase gene expression (Figure S6) and enzyme activity (Figure 4B) remained elevated in diploids and triploids following multiple stress exposure. The mitochondrial enzyme citrate synthase (CS) catalyzes the first step of the TCA cycle (Ciccarone et al., 2017) and is correlated with respiration rates within marine invertebrates (Dahlhoff et al., 2002; García-Esquivel et al., 2001, 2002). Sustained CS activity within triploids as they attempt to decrease metabolic demand could constitute a substantial energetic burden. However, it remains unclear whether the observed CS activity following stress exposure is attributable to ploidy, as studies of CS activity within other taxa have produced mixed results. For example, triploid zebrafish displayed the same CS enzyme activity following temperature stress (van de Pol et al., 2021), while triploid white sturgeon displayed a marked decrease with respect to diploids (Leal et al., 2019). Nevertheless, meaningful differences in standard metabolic rate were observed across ploidy within this study, with triploids decreasing and diploids increasing oxygen consumption following multiple stress exposure (Figure 3F). This difference suggests that triploids have a lower metabolic scope than diploids and is consistent with triploids exhibiting a lower critical thermal temperature. A reduction in aerobic scope under high temperature has been proposed to contribute to triploid mortality within Atlantic salmon, particularly under decreased oxygen saturation (Sambraus et al., 2017). The potential for polyploidy to prevent or slow the transition from aerobic to anaerobic metabolism through the dysregulation of important genes that function in ATP generation and turnover is an interesting hypothesis that warrants further study.

### 4.6. Conclusion

Our results contribute to a growing body of work that suggests that polyploidy negatively impacts environmental stress tolerance through the dysregulation of key metabolic processes, the consequences of which are evident following extreme climate events such as MHWs. A comparison of mortality rates across ploidy within this study supports the prevailing view that temperature stress alone is unlikely to motivate triploid mortality events observed in aquaculture. Instead, we hypothesize triploid mortality arises from ploidy-specific differences in energy allocation that manifest within multiple stressor scenarios (T.-C. F. Pan et al., 2015), resulting in energetic-limitations within critical biological processes related to stress tolerance (Sokolova, 2013). Our results provide evidence that polyploidy sufficiently alters the transcriptional response of Pacific oysters to environmental stress, resulting in downstream shifts in physiological tolerance limits that may be detrimental to organismal survival.

## Contributions

MNG, MG, BV and SR conceived of the study, analyzed data, and wrote the manuscript. MNG oversaw the completion of hatchery experiments, conducted respirometry measurements, sampled tissue, performed RNA extractions, and analyzed sequencing results. BV and OC assisted with tissue sampling during hatchery experiments. OC completed the citrate synthase assay. DL completed the histology analysis. MM completed the ATPase assay.

## Supporting information

table_S6-3n-SS-DEG

table_S7-2n-MS-DEG

table_S8-3n-MS-DEG

table_S5-2n-SS-DEG

## Acknowledgements

We are grateful to Dr. Ariana Huffmyer’s assistance with tissue sampling, Dr. Hollie Putnam for sharing equipment, Kelly Hudkins and the University of Washington Histopathology laboratory for assistance with histological sectioning and staining, and the support of the hatchery team at the Jamestown S’Klallam Point Whitney Shellfish Hatchery led by Matthew Henderson and Nathan Tsao.

## Data Availability

Raw sequencing data files are stored on NCIBI under the BioProject ID 913164. Data and code used for analyses can be found at https://doi.org/10.5281/zenodo.7693092.

## 6. Supplemental Information

**Figure S1.**
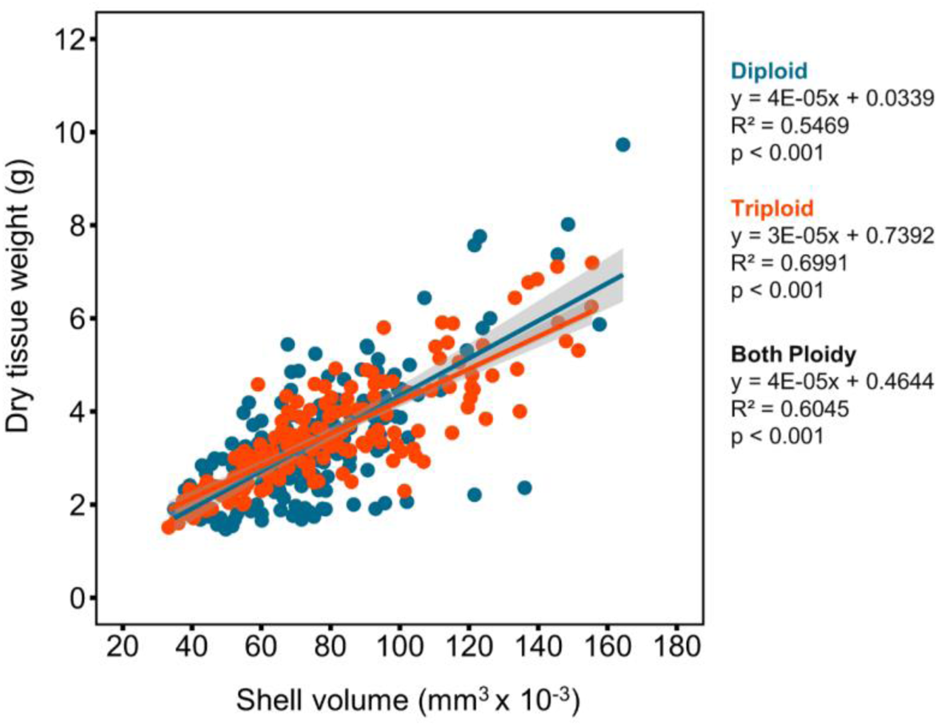
Relationship between measured shell volume (mm^3^) and whole-body dry tissue weight (g) for diploid (blue) and triploid (orange) oysters used within experiments. A linear regression model fit across both ploidy was used to estimate dry tissue weight from measured shell volume when the determination of dry tissue weight for an individual was not possible.

**Figure S2.**
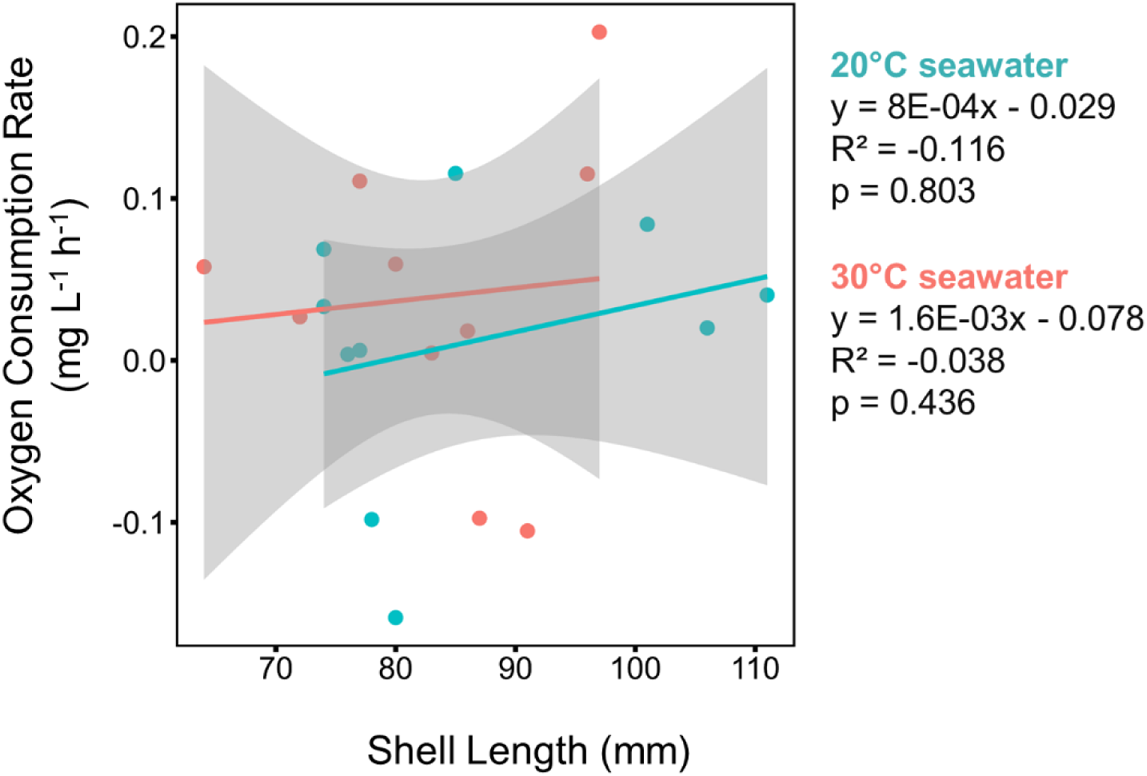
Relationship between oxygen consumption rate and size of oyster shell used within blank respirometry chambers.

**Figure S3.**
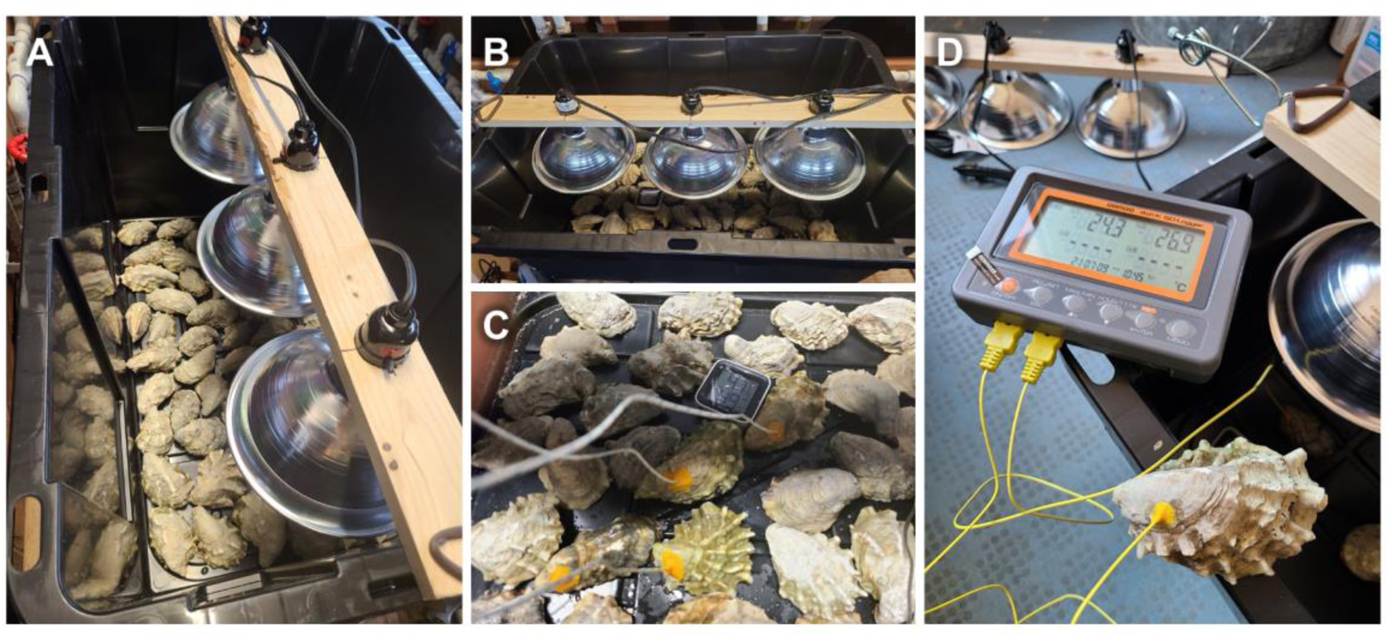
Desiccation chamber (A-B) used to apply secondary stress treatment on day 2 within the multiple stress treatment (2n-MS and 3n-MS treatment groups). The air temperature within each chamber simulated conditions present within the intertidal zone at low tide during a heat wave; the conditions experienced by oysters within chambers is presented in Figure 1B. The impact of chamber air temperature on the internal temperature of oysters was assessed by drilling a hole in the top valve of a subset of oyster and inserting a thermocouple into the stomach (C-D); the maximum internal oyster temperature as a function of shell volume is presented in Figure 1C.

**Figure S4.**
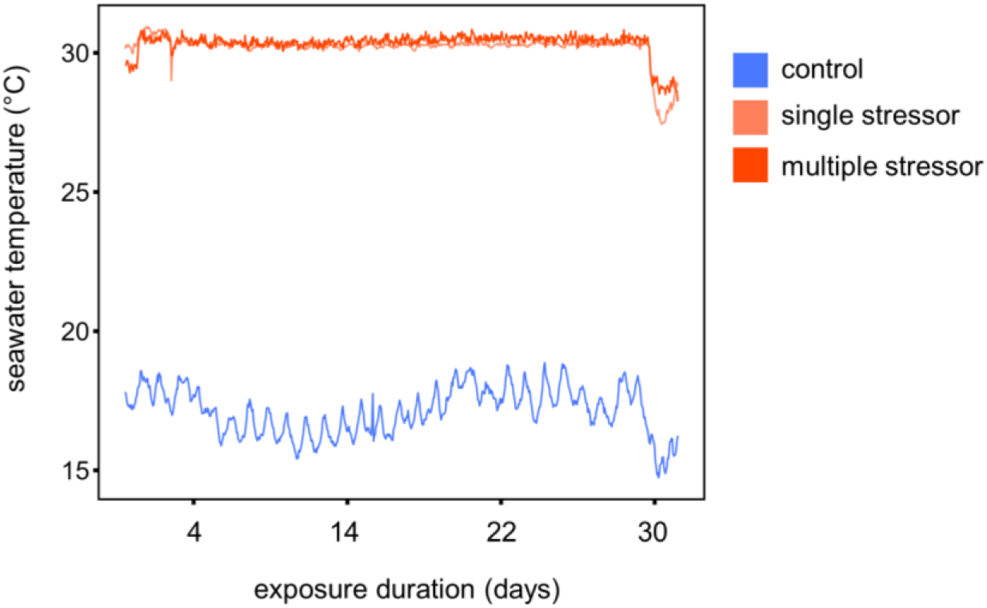
Seawater temperature within each treatment condition (control, single stressor, multiple stressor).

**Figure S5.**
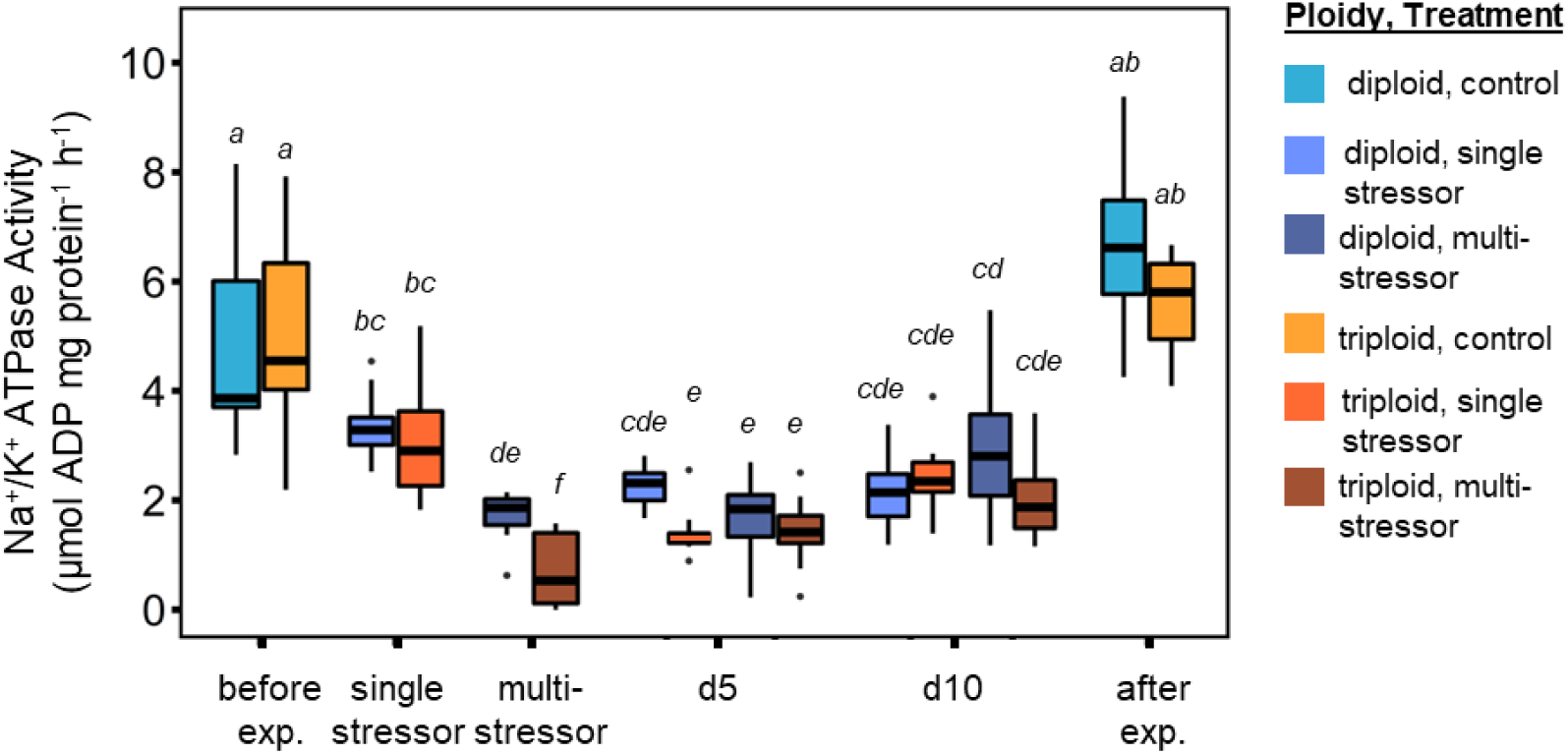
The impact of ploidy, stress type, and exposure duration (days) on Na^+^/K^+^ ATPase (A; n=10-13 per treatment; 1, 5, and 10 days after stress) and citrate synthase (B; n=9-12 per treatment; 1 day after stress) enzyme activity of adult Pacific oysters. All oysters were acclimated to hatchery conditions (20d, 20°C) before inclusion in either the control (20°C), single stressor (SW temp = 30°C), or multi-stressor (SW temp = 30°C; aerial exposure = 44°C for 4h) treatment. Italicized letters represent the result of post-hoc Tukey HSD comparisons across groups; boxplots that share a letter are not significantly different. All enzyme activity was measured within the ctenidium.

**Figure S6.**
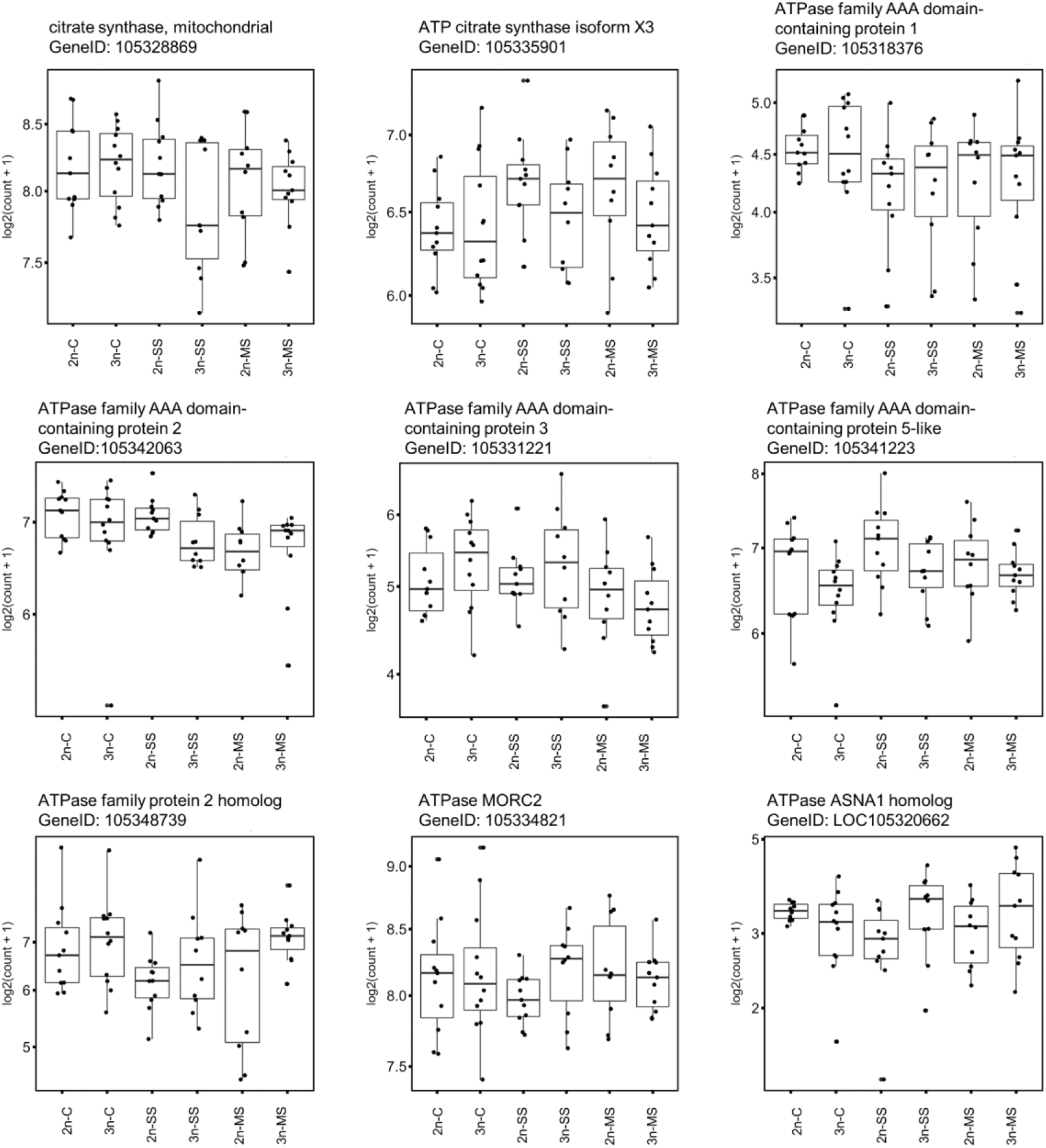
Expression of citrate synthase and ATPase genes across all treatments

**Table S1.** Mean (±SD) shell length, shell width, shell height, and dry tissue weight from diploid (2n) and triploid (3n) oysters upon arrival at the hatchery (baseline condition) and within the control, single stressor, and multiple stressor treatments on day 30 (end of experiment).

| Treatment | Ploidy | Shell Length (mm) | Shell Width (mm) | Shell Height (mm) | Dry tissue weight (g) |
| --- | --- | --- | --- | --- | --- |
| Baseline condition | 2n | 90.0 ± 7.8 | 55.6 ± 4.8 | 32.3 ± 3.6 | 3.9 ± 0.7 |
| | 3n | 76.8 ± 10.1 | 58.3 ± 8.3 | 29.8 ± 4.1 | 3.3 ± 0.9 |
| Treatment control | 2n | 87.8 ± 9.4 | 54.5 ± 7.1 | 30.7 ± 4.5 | 3.6 ± 1.0 |
| | 3n | 78.6 ± 10.0 | 56.5 ± 7.5 | 31.6 ± 5.0 | 3.5 ± 1.0 |
| Single stressor | 2n | 85.9 ± 8.4 | 52.9 ± 6.4 | 29.6 ± 3.6 | 3.3 ± 0.7 |
| | 3n | 83.5 ± 8.5 | 58.8 ± 5.3 | 31.8 ± 3.4 | 3.8 ± 0.9 |
| Multiple stressor | 2n | 86.2 ± 9.4 | 52.6 ± 6.5 | 30.0 ± 4.0 | 3.3 ± 0.8 |
| | 3n | 81.9 ± 9.0 | 58.6 ± 8.8 | 31.9 ± 4.7 | 3.8 ± 1.2 |

**Table S2.** Two-way analysis of variance (ANOVA) results comparing the effect of ploidy, stress treatment (control, single stressor, or multiple stressor), and their interaction on oyster shell length, shell height, shell width, and dry tissue weight.

| Factor | Source | DF | SS | F | p |
| --- | --- | --- | --- | --- | --- |
| Shell Length | ploidy | 1 | 6,493 | 75.7 | <0.001 |
|  | treatment | 3 | 204 | 0.79 | 0.50 |
|  | ploidy:treatment | 3 | 1,635 | 6.36 | <0.001 |
|  | residuals | 602 | 51,613 | - | - |
| Shell Height | ploidy | 1 | 10.9 | 11.1 | <0.001 |
|  | treatment | 3 | 0.4 | 0.14 | 0.94 |
|  | ploidy:treatment | 3 | 11.9 | 4.06 | 0.007 |
|  | residuals | 602 | 586 | - | - |
| Shell Width | ploidy | 1 | 10.9 | 11.1 | <0.001 |
|  | treatment | 3 | 0.4 | 0.14 | 0.94 |
|  | ploidy:treatment | 3 | 10.2 | 3.76 | 0.011 |
|  | residuals | 602 | 545 | - | - |
| Dry Tissue Weight | ploidy | 1 | 2.4 | 2.47 | 0.117 |
|  | treatment | 3 | 0.4 | 0.14 | 0.94 |
|  | ploidy:treatment | 3 | 19.1 | 6.54 | <0.001 |
|  | residuals | 602 | 587 | - | - |

**Table S3.**
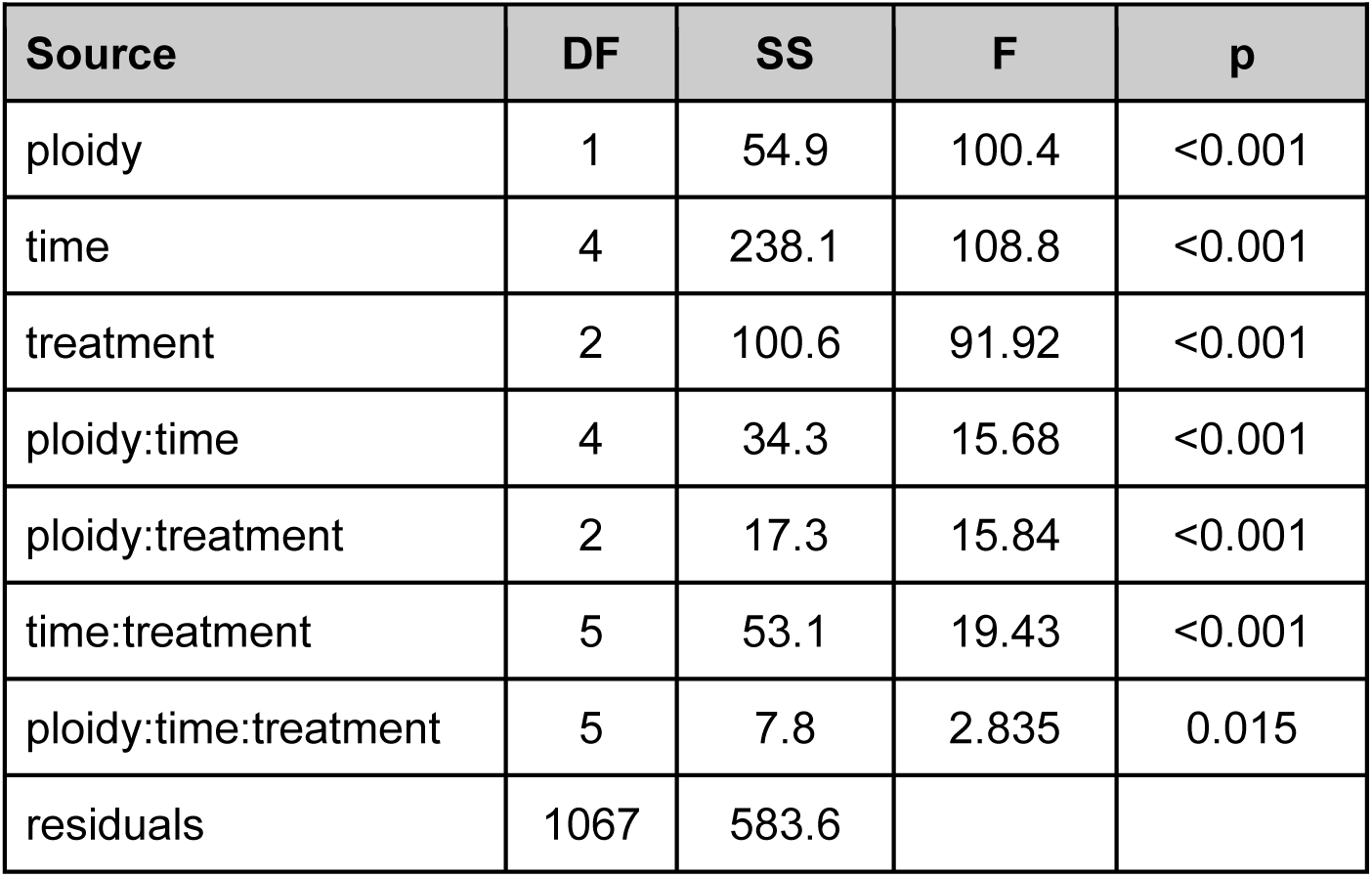
Repeated measures analysis of covariance (ANCOVA) results comparing the effect of ploidy (diploid, triploid), stress treatment (control, single stressor, or multiple stressor), exposure duration (days; −10, 1, 2, 5, 10), and their interaction on oyster metabolic rate.

**Table S4.**
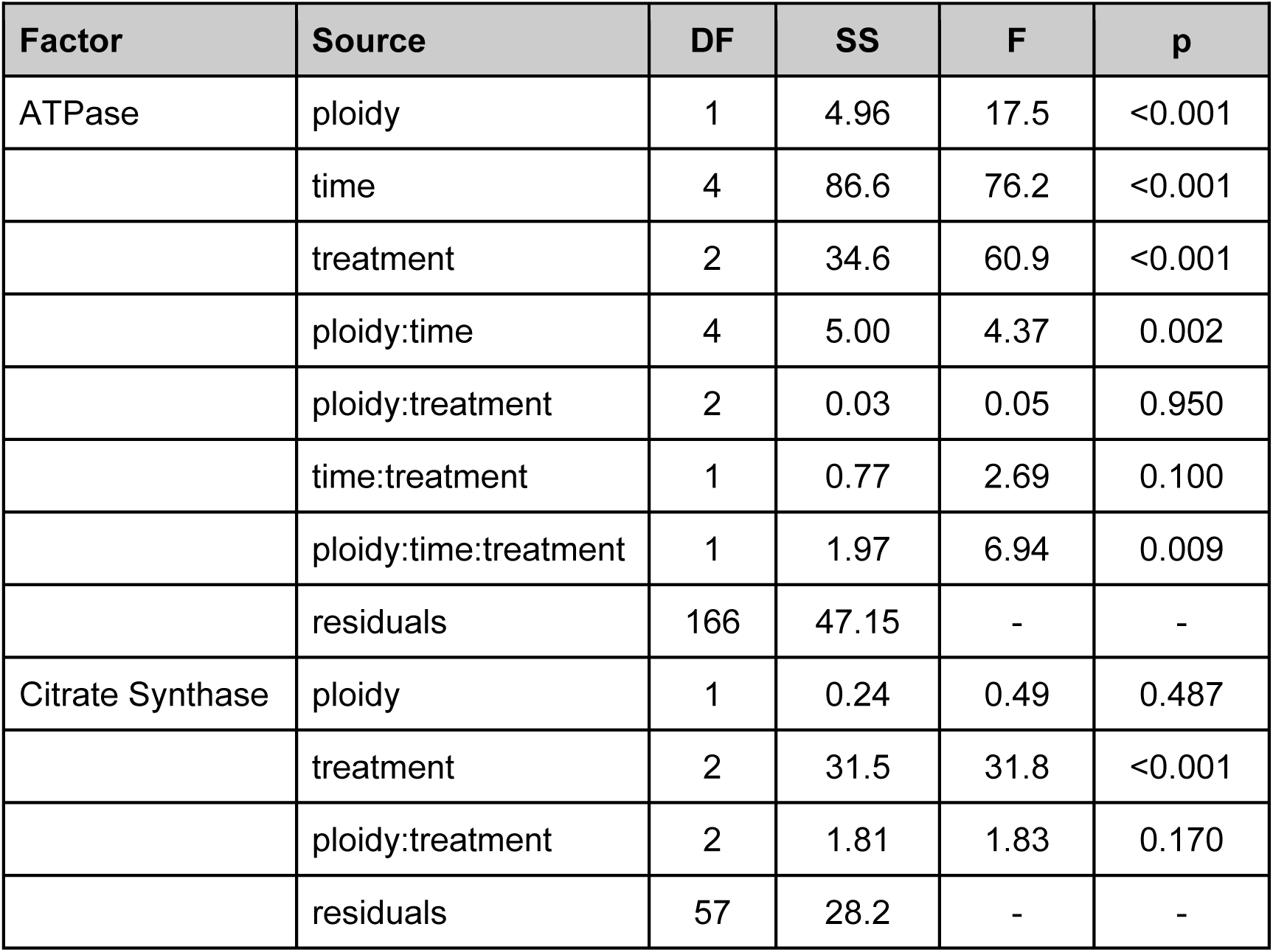
Analysis of covariance (ANCOVA) results comparing the effect of ploidy, stress treatment (control, single stressor, or multiple stressor), exposure duration (days), and their interaction on Na^+^/K^+^ ATPase and citrate synthase activity within the ctenidia.

**Table S5.** Significant differentially expressed genes (DEG) within diploid (2n) Pacific oyster following single stress exposure with respect to the diploid control.

< see table_S5-2n-SS-DEG.csv>

**Table S6.** Significant differentially expressed genes (DEG) within triploid (3n) Pacific oyster following single stress exposure with respect to the triploid control.

< see table_S6-3n-SS-DEG.csv>

**Table S7.** Significant differentially expressed genes (DEG) within diploid (2n) Pacific oyster following multiple stress exposure with respect to the diploid control.

< see table_S7-2n-MS-DEG.csv>

**Table S8.** Significant differentially expressed genes (DEG) within triploid (3n) Pacific oyster following multiple stress exposure with respect to the triploid control.

< see table_S8-3n-MS-DEG.csv>

**Table S9.** Ploidy-specific gene ontology terms (GOterms) that were significantly enriched within diploid (2n) Pacific oyster following single stress exposure.

< see table_S9-2n-SS-GO.csv>

**Table S10.** Ploidy-specific gene ontology terms (GOterms) that were significantly enriched within triploid (3n) Pacific oyster following single stress exposure.

< see table_S10-3n-SS-GO.csv>

**Table S11.** Ploidy-specific gene ontology terms (GOterms) that were significantly enriched within diploid (2n) Pacific oyster following multiple stress exposure.

< see table_S11-2n-MS-GO.csv>

**Table S12.** Ploidy-specific gene ontology terms (GOterms) that were significantly enriched within triploid (3n) Pacific oyster following multiple stress exposure.

< see table_S12-3n-MS-GO.csv>

